# In Other Words: Learning When to Trust Generated Protein Function Descriptions

**DOI:** 10.64898/2026.10.15.757758

**Authors:** Tao Li, R. Prabakaran, Rund Tawfiq, Frederic Guintu, Carl Yang, Yana Bromberg

## Abstract

Generative protein-to-text models write function descriptions in free-form English but give no indication of which to believe. Indeed, free text is hard to evaluate. We introduce a large language model judge of equivalence of predicted vs. “true” function descriptions, validated against blinded expert curation. We also present ProtTale, our sequence-to-text model, that learns from the judge’s verdicts to estimate its own reliability, *pJay*. On unseen Swiss-Prot enzymes, ProtTale and three other leading methods tie on accuracy but are right about different proteins. Here, ProtTale’s unique correct predictions outnumber all other methods’ combined. Scored from sequence alone, pJay is a plug-in filter, raising all models’ performance. On short proteins, it raises precision for every tested method nearly twofold. We argue that generative annotation becomes useful not by being right more often, but by knowing when it can be trusted—a standard that every generative model in biology should meet.

---

A typical cell runs on proteins, yet our ability to say what any one of them does remains poor [1–4]. Manual curation covers 0.38% of known proteins (UniProtKB release 2026 02); everything else is, at best, computationally annotated [5], or not annotated at all.

The reliable route has long been homology, i.e. sequence alignment, structure comparison, and profile methods that treat shared ancestry as a proxy for shared function [6–11]. Domains and families serve as reusable functional units [12–14], and interaction context adds an orthogonal axis [15, 16]. These methods work well when the new protein and the annotated one are truly similar, but they cannot describe what the reference collection does not already contain. However, multiple constraints limit both the accuracy and coverage of automated *in silico* annotation [4]. For example (1) divergent evolution can preserve sequence and/or structure while shifting functional scope, regulation, or context of action [17], (2) convergent evolution can produce similar functions with minimal similarity [18], and (3) existing experimental biases limit annotations to a narrow slice of the protein space [19].

Protein language models (pLMs) were supposed to help, and in part they have. Their embeddings seem to capture remote homology, biochemical constraints, and aspects of structure [20, 21], and deep models built on them assign Gene Ontology terms well [22–25]. Proteins far from the curated space, however, remain hard [3, 4, 26]. What is left is a “functionally dark” protein space, full of proteins that may matter medically or environmentally but that nobody can describe [27, 28]. Note that darkness is not the only problem. Proteins of the *transparent proteome* have detectable, sometimes annotated, homologs that share no common function, which leaves the protein itself just as unresolved [29].

A deeper constraint is vocabulary. Gene Ontology terms, Enzyme Commission classes, and pathway catalogs are discrete bins, built to keep annotation consistent across organisms [30–32]. Protein function is not discrete. One family can do different things depending on context, partners, and conditions, and fixed labels capture this only partially. Recent protein-to-text models sidestep the bins by writing descriptions in natural language [33, 34]. Expressiveness, however, comes at a price: free text reads well whether or not it is right, it resists quantification downstream, and no standard metric can tell whether two descriptions mean the same thing. Controlled vocabularies thus remain essential [4], and here we use ontology extraction as a lens on free text.

Here, we present ProtTale, which asks the question its predecessors do not: not only *what does this protein do?* but *should you believe the answer?* We supervise a representation module with machine-inferred protein similarity (MIPS), i.e. labels read off embedding geometry over millions of largely uncharacterized sequences rather than from a curator [35, 36], so that functionally similar proteins land near one another. We then train the model to emit its own reliability estimate, in the spirit of pLDDT in structure prediction [37], so that every annotation arrives with a statement of how much to trust it. We benchmark against current protein-to-text and multimodal methods [33, 38, 39], score functional equivalence with an LLM-as-judge protocol rather than surface overlap, and validate with blinded expert curation. Our aim is to shrink both the dark and the transparent regions of protein space, turning accumulated sequence into knowledge one can use [27–29].

## Results

### Embedding proximity is a usable, if imperfect, proxy for shared function

Protein language models (pLMs) seem to encode evolutionary, physicochemical, and structural constraints in the geometry of their latent space. We use this machine-learned biology as an explicit supervisor, i.e. we turn embedding proximity into training labels. However, we first asked whether proximity is a defensible proxy for shared function at all.

Using the ProtT5 protein language model (pLM), we embedded 5.2 million UniRef90 proteins (that have ≥10 homologs) and performed large-scale clustering over the 5.5 billion edges connecting proteins with similar embeddings (cosine similarity *>* 0.77); this yielded 1.2 million communities (Methods). Although embedding proximity does not guarantee sequence, structure, or function similarity, our evaluation of 1,000 pairs of annotated Swiss-Prot proteins sharing the same cluster (244 clusters considered) showed a mean pairwise sequence identity of 37.5% (95% CI: 36.5–38.6%), 58.8% (95% CI: 55.7–61.8%) had structures with TM-score ≥0.7, and 63.1% (95% CI: 60.1–66.0%) shared identical EC numbers at the fourth EC level, providing empirical support for the use of embedding proximity as a proxy for similarity (Methods; Supplementary Note 1).

These 1.2 million clusters define our Machine-Inferred Protein Similarity (MIPS) labels. Pairs from the same cluster are labelled similar (positive) and pairs from different clusters dissimilar (negative). Proteins in different clusters may still share function, so a negative label means “not detectably similar in embedding space” rather than functionally different.

ProtTale trains in two stages. Stage 1 aligns sequence- and text-derived representations using our MIPS-labelled protein pairs (Extended Data Fig. 1A), where the pairing comes from embedding geometry but each pair is supervised by both proteins’ curated Swiss-Prot descriptions. Stage 2 then trains the generator on Swiss-Prot proteins paired with their curated FUNCTION descriptions (Extended Data Fig. 1B). Table 1 gives the sizes and roles of every data set split used here. Splits are defined at the level of CD-HIT [40] clusters built at 40% sequence identity, so that no protein in one split has a close homolog in another, and the unseen set is drawn from clusters that entered Swiss-Prot after 2022 (Methods).

**Table 1:** Sizes and roles of each dataset split. Stage 1 pairs are balanced, with equal numbers of positive (within-cluster) and negative (across-cluster) pairs per split. Dashes mark splits not used for a given purpose: the unseen set contributes no Stage 1 pairs, and the reliability head skips the training split.

| Split | Dataset size |  | Role in training |  |
| --- | --- | --- | --- | --- |
|  | Stage 1 pairs | Stage 2 proteins | Generator | Reliability head |
| Training | 119,546 | 207,197 | Training | – |
| Validation | 388 | 1,000 | Validation | Training |
| Testing | 534 | 1,000 | – | Validation |
| Unseen | – | 1,031 | Benchmark | Benchmark |

### Mapping free text to GO terms adds noise on the scale of the method differences

Function prediction has traditionally been scored on recall of Gene Ontology (GO) or Enzyme Commission (EC) term sets [4]. Free-text predictions instead need a metric of functional equivalence. BLEU [41] and ROUGE [42] reward *n*-gram overlap, and BERTScore [43] returns a semantic score that is hard to interpret for specific biological claims. The obvious alternative is to map text back to an ontology, so we asked whether this mapping carries a reliable signal (Methods).

We collected curated-text function descriptions for the 1,031 **unseen** proteins, none of which appears in the training data of any method compared here (Table 1; Methods). We used Claude Opus 4.7 (Supplementary Note 2) to extract their GO terms and compared these against the ground-truth Molecular Function (MF) annotations of each protein using Wang semantic similarity [44] without ancestor expansion. To anchor the absolute scale of the metric, we computed random baselines by shuffling the “true protein GO terms” vs. “extracted GO term assignment” pairs.

We then predicted function descriptions for these proteins with ProtTale and four published protein-to-text methods spanning the architectural space of the field, ESM2Text, Prot2Text, ProtT3 and Protein2Text, all run from their released checkpoints without retraining (Methods), and evaluated the extracted GO term sets.

Note that during the extraction process, we observed a practical issue: some calls (e.g., “Extended-spectrum beta-lactamase (ESBL) which confers resistance to penicillin, as well as first, second, and third-generation cephalosporins. Has cefotaxime-hydrolyzing activity.”) triggered Claude’s content-safety refusals and no returned GO terms. These count as empty extractions, reported per method and excluded from the similarity averages for all methods alike.

**Fig. 1:**
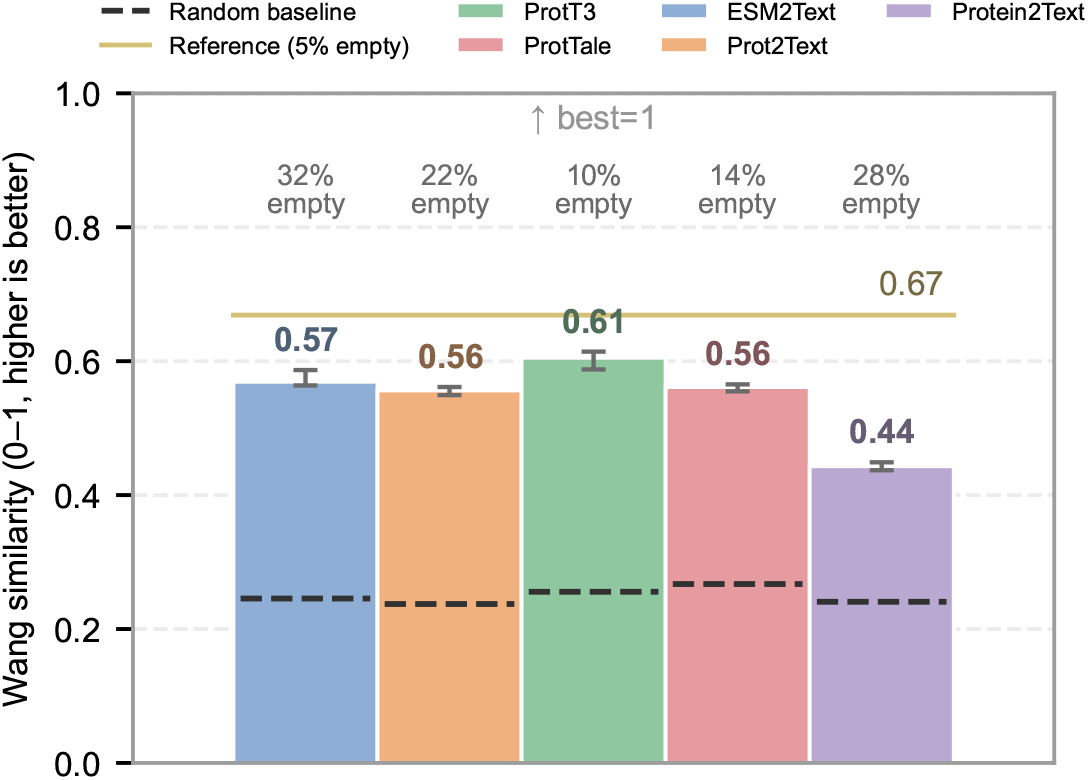
Extracting GO terms from free text imposes a low performance ceiling and flattens differences between methods. Wang similarity (without ancestor expansion) between GO terms extracted from predicted free-text annotations and ground-truth MF GO terms on the unseen protein set. Bars show means over five runs and error bars span the minimum and maximum across runs, and percentages above bars give the empty-extraction rate, i.e. the fraction of inputs yielding no GO terms. The solid olive line marks the curated-text reference, obtained by applying the same extraction to curated Swiss-Prot descriptions; the dashed black line marks a random baseline in which extracted and true GO term sets are shuffled between proteins.

Every method performed above the random baseline but below the curated-text ceiling (Fig. 1). Empty-extraction (no GO terms retrieved from text) rates ranged from 10% (ProtT3) to 32% (ESM2Text). The four leading methods clustered close to the curated-text ceiling. Swiss-Prot descriptions are correct by construction, yet after GO extraction they reach a Wang similarity of only 0.67 against the entries’ own GO terms, so the missing third is lost in the mapping itself. This loss is nearly seven times the spread among the four leading methods (Prot2Text 0.556, ProtTale 0.561, ESM2Text 0.569, ProtT3 0.605), with Protein2Text at 0.443.

Similar conclusions held for (1) Wang similarity with ancestor expansion, (2) Jaccard similarity [45], and (3) information accretion metrics [46] (Extended Data Fig. 2; Supplementary Note 3).

The limitations of this evaluation approach motivated us to forgo ontology mapping and directly judge predictions at the free-text level.

### LLM-as-judge is precise, conservative, and reproducible

To compare two free-text descriptions directly, we adopted the LLM-as-judge paradigm [47], in which a large language model scores candidate outputs in place of a human. We introduce a domain-specific prompt that returns one of four labels: *Yes* for a confident match, *No* for a confident mismatch, *Maybe* for a specific but inconclusive comparison, and *Vague* for outputs too underspecified to evaluate, returned by the prompt as scores of 1, 0, 0.5 and −1 (Methods, prompt in Supplementary Note 2). By analogy (not part of the prompt), *Yes* compares two Toyota Camrys, *Maybe* a Camry with a Honda Civic, *No* a Camry with a cheesecake, and *Vague* a Camry with a generic “vehicle”.

To evaluate judge-model accuracy, we used the 534 held-out pairs of the **Stage 1 test set**, MIPS-labelled pairs of Swiss-Prot proteins (Table 1; Methods), with their curated descriptions. Same-cluster pairs are expected to share function and different-cluster pairs to be enriched for different functions, so an uninformative judge would assign *Yes* equally to both.

In testing, only ∼4% of pairs (20 of 534) were labelled *Yes*, likely because Swiss-Prot descriptions often combine several functional statements and the prompt assigns *No* whenever any of them lacks a counterpart. Of these 20, 18 were same-cluster pairs (95% Wilson CI: 69.9–97.2%; Fisher’s exact test, *P* = 3.2 × 10*^−^*^4^; Extended Data Fig. 3A), against 50% expected for an uninformative judge. By contrast, the 490 *No* labels split almost evenly between same-cluster (231) and different-cluster (259) pairs. That is, the *No* label is much less informative than the *Yes* signal.

To further evaluate judge-model consistency, we performed five independent runs on every method’s unseen-set predictions (Methods). Agreement between runs was 75–81%, and Cohen’s *κ* ranged from 0.61 to 0.73 (Extended Data Fig. 3B). In standard interpretation [48], *κ* ≥ 0.6 indicates substantial agreement. The judgement stability for all methods met or exceeded this threshold. We treated a prediction as *Yes* when at least three of the five runs said so, and use that majority throughout.

These results indicate that the LLM-as-judge is highly precise, if deliberately conservative. As such, *Yes* rates throughout this work should be read as lower bounds on functional agreement rather than as accuracy.

### Free-text predictors show similar accuracy but make orthogonal predictions

The central question for any predictor is how it fares far from the homology neighborhoods of the curated reference. The unseen set is such a case: no protein in it shares a 40% identity cluster with anything used in training, 47% have no homolog at that threshold anywhere in Swiss-Prot, and every protein carries an EC number, so the benchmark measures enzyme function specifically. Other methods were checked for exact training overlap with the unseen set sequences, which guarantees that none of these proteins leaked into their training data (Methods).

Altogether, at least one method correctly described 450 of the 1,031 unseen proteins, i.e. 44%, leaving 581 that no method described correctly (Extended Data Fig. 4). Four of the five methods reached comparable *Yes* rates of 24.9–26.5%, while Protein2Text reached only 6.2% (Fig. 2A). All six pairwise McNemar tests among the four leading methods gave *P*_holm_ = 1.00, whereas Protein2Text differed from each at *P*_holm_ *<* 10*^−^*^34^.

The methods, however, were right about different proteins (Fig. 2B, Extended Data Fig. 4). Of the three methods shown, 109 proteins were correctly identified by all of them; ProtTale uniquely identified 79, compared with 59 for Prot2Text and 34 for ProtT3. In the five-method comparison, only 17 proteins were described correctly by every method, while ProtTale uniquely recovered 66, i.e. more than the other four methods’ unique sets put together (65). Pairwise Cohen’s *κ* on the four-label predictions showed limited agreement across methods (Fig. 2C; method-pair *κ* = 0.10–0.34, except for Prot2Text vs. ESM2Text, *κ* = 0.60).

These differences reflect systematic biases in the models’ ability to predict function (Fig. 2D). For ESM2Text, Prot2Text and ProtT3, the number of the protein’s known sequence-similar entries significantly influences how accurately the its function can be predicted. ESM2Text and Prot2Text were also sensitive to the level of similarity of the best structural match available in the PDB, indicating the influence of structure-driven training of their foundation models. ProtT3’s accuracy was, on the other hand, influenced by the number of good (TM score ≥ .7) structural matches [49, 50] in PDB[51]. Curiously for ProtT3, the trend is that larger structural family sizes indicate worse function prediction performance. This may indicate the fine-tuned model’s effort to capture features explicitly relevant to function, not structure, prediction. ProtTale’s prediction success, however, was not influenced by the sequence or structure signals in protein representations. These results clearly show that different architectures and training data induce distinct biases, with each method covering a different region of protein space. Note, however, that ProtTale captures the largest unique subset of the methods compared here.

### The reliability head scores the protein, not the description

The fact that models land in a narrow space, only 450 of the 1,031 proteins across all methods, suggests that they sit near a shared limit for “understanding” of unseen proteins. The question is thus not “Which method is best?” but “For any method, which prediction can I trust?” Note that this matters most to the experimentalist, who has to spend bench time on whichever annotation they believe.

We framed reliability as binary, *Yes* against all other labels, and trained a head on frozen ProtTale features, the mean-pooled encoder and Q-Former outputs, to predict the probability that the judge would call ProtTale’s description a match (Methods). We call this score *p*_JAY_, the predicted probability of a Judge-Assigned *Yes*. Because the head reads the sequence representation only, on unseen proteins and on other methods’ predictions *p*_JAY_ is a statement about the protein, not about any particular description.

**Fig. 2:**
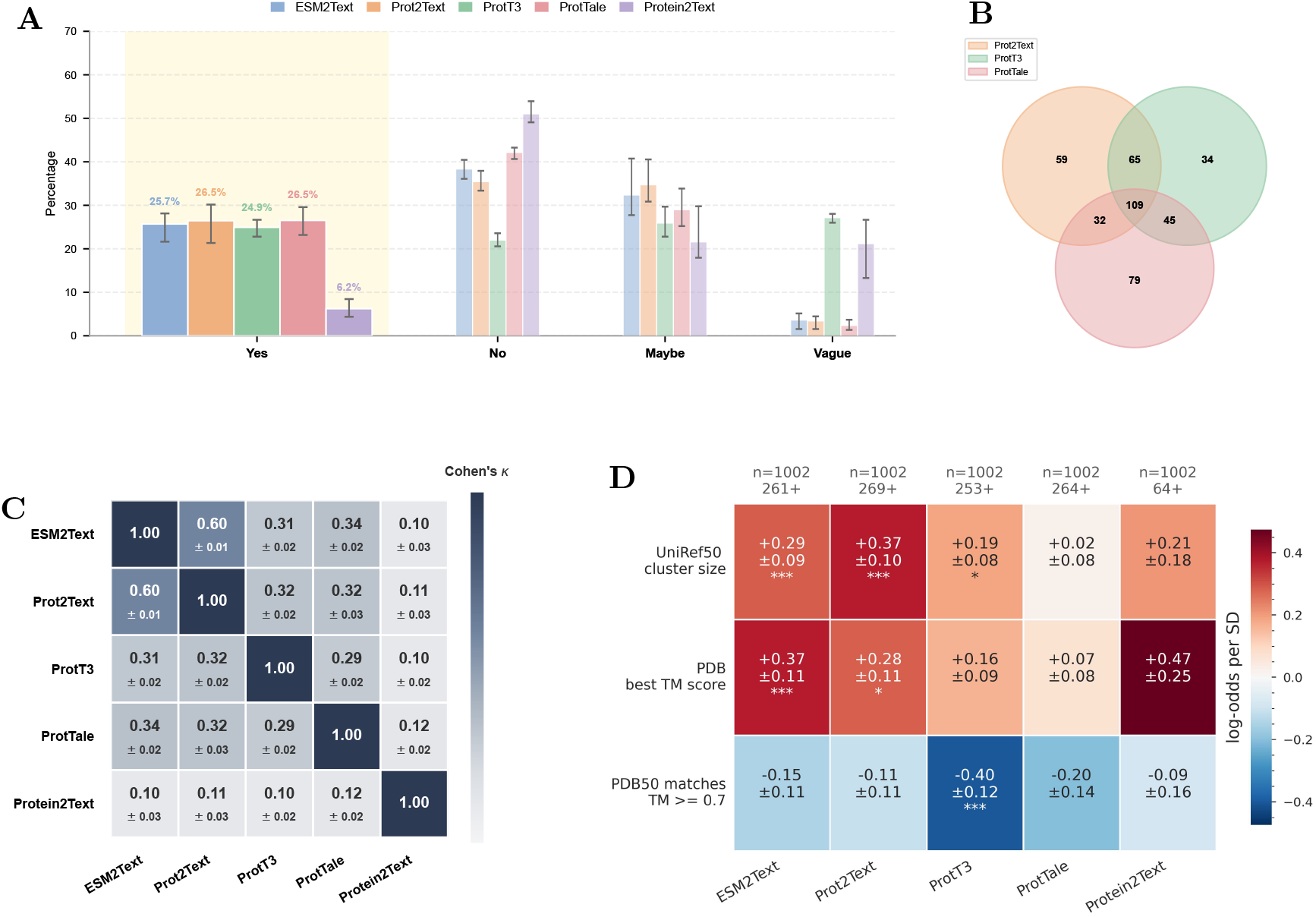
The four leading protein sequence-to-text methods reach similar accuracy but succeed on different proteins. **(A)** Distribution of LLM-as-judge labels per method, with *Yes* shaded; Protein2Text is the sole outlier. Bars are means over the five judge runs; all counts elsewhere use the five-run majority. **(B)** Overlap of *Yes* predictions across three methods. ESM2Text is omitted because it shares a backbone with Prot2Text, which additionally incorporates predicted structure. Protein2Text is omitted because its 6.2% *Yes* rate produces too few correct predictions to populate the diagram legibly. The five-method version is in Extended Data Fig. 4. **(C)** Pairwise Cohen’s *κ* between methods on the four-label predictions, with standard error across runs; low values indicate that methods agree little beyond chance. **(D)** The extent to which known function annotation contributors, i.e. family size and best match in PDB, influence a method’s function prediction. Logistic-regression (LR) derived log-odds scores are computed on the unseen set for: UniRef50 cluster size (sequence family), best TM score vs. PDB100 (structural similarity to experimentally determined structures), and the number of matches in PDB50 with at least TM score of 0.7 (structure family). For each method, an LR model was trained to predict whether that method’s functional annotation was judged correct by the LLM-Judge. Mean log odds, with standard error, are reported for each feature, averaged over five LLM-Judge replicates with 10-fold cross-validation. Significance was assessed by Wald tests on the fitted coefficients: *^∗^P <* 0.05, *^∗∗^P <* 0.01, *^∗∗∗^P <* 0.001. Note analysis on Protein2Text prediction did not result in statistically significant claims due to the large variations across runs.

To anchor where signal begins, we used shuffled amino acid sequences, i.e. residue permutations of the same unseen proteins. Real proteins extend a long right tail in the *p*_JAY_ distribution that shuffled sequences never reach; a Fisher exact sweep places the most significant threshold for separation at *τ ^∗^* = 0.537 (Extended Data Fig. 5; Supplementary Note 4). We thus adopted *p*_JAY_ *>* 0.537 as the discrete reliability filter for trusted predictions. Of the 1,031 unseen proteins, 201 (19.5%) scored above this cutoff, while the remaining 830 (80.5%) were deemed too difficult to annotate.

ProtTale truncates inputs at 1,024 residues (Methods), which covers 92% of viral, 94% of eukaryotic and 99% of bacterial Swiss-Prot proteins. We therefore report a short (≤ 1,024 residues, *n* = 852) and a long (*n* = 179) subset separately, long-subset predictions resting on the N-terminal 1,024 residues alone.

Generated descriptions are capped at 128 tokens, and 233 of the 1,031 unseen predictions (22.6%) reach the cap without emitting an end-of-sequence token (Methods).

Applying the reliability filter raises the precision (*Yes* rate) of every method on short proteins and of all but ProtT3 on long ones (Fig. 3A,B), and raises Wang similarity for ProtT3 and ProtTale (Extended Data Fig. 6A,B). Because the head reads only the protein, it scores how *predictable* a protein’s function is rather than whether a given description is correct; that the filter transfers to methods whose output it never saw supports this reading. The same logic underlies RNS, which scores whether a protein embedding is trustworthy without reference to any downstream prediction made from it [52].

Some functions are intrinsically easier to infer from sequence, because they are “simpler” to read, more uniquely defined, or more common in the known functional space on which all models are trained. Family size supports this reading: reliability rises with the size of a protein’s 40% identity cluster, from a median of 0.33 for the 47% of unseen proteins with no homolog at all to 0.37 for those in clusters of twenty or more (Mann–Whitney, *P* = 0.03 for singletons against the rest; Supplementary Data 2). Note that the model saw none of those relatives, since all other cluster members were withheld from every split, so this reflects how well populated a functional space is rather than exposure during training. For short sequences the enrichment was significant across the four leading methods (Fisher exact, *P*_holm_ ≤ 4 × 10*^−^*^4^). On this short subset ProtTale rose from 25.8% to 45.9% precision (from 0.56 to 0.61 Wang similarity), and ProtT3 ended up at a similar level (46.4% precision, 0.64 Wang similarity; McNemar, *P* = 0.64).

For long sequences, ProtTale performed best numerically, with a 32.1% *Yes* rate and 0.59 Wang similarity, although the small sample (*n* = 48 of 179, after filtering) limits statistical inference. Since proteins longer than 1,200 residues contributed no Stage 1 pairs (Methods), ProtTale’s lead on long proteins does not come from Stage 1. Indeed, retraining ProtTale without Stage 1 halves the *Yes* rate on short proteins, from 25.8% to 12.7% (McNemar, *P* = 7.7 × 10*^−^*^16^), leaving long proteins unchanged (29.8% versus 29.1%, *P* = 0.60; Extended Data Fig. 7), as expected. MIPS-supervised alignment is thus what makes ProtTale competitive where Stage 1 pairs exist. The continuous *p*_JAY_ score further lets users tune precision vs. coverage. At *p*_JAY_ *>* 0.75, 20 of the 1,031 predictions remain and ProtTale reaches 66% precision, against 61% for ProtT3 and about 50% for the other two. Precision climbs to about 90% at the highest cutoffs, though few predictions remain there (*n* = 2; Fig. 3C). Note that neither *p*_JAY_ itself nor the set of proteins retained at each *p*_JAY_ threshold (Fig. 3d) could be predicted from the usual function prediction directions, i.e. available sequence- or structure-similar entries in the known protein space. This indicates that our reliability score improves precision of function prediction without simply tracking homology to already-characterized proteins.

### Reliability filtering helps in every taxonomic group

The unseen set is dominated by Eukaryotes (604 of 1,031; 59%), so the results above largely describe them. We asked whether they also hold for Bacteria, Archaea and Viruses, excluding subsets with fewer than three proteins left after filtering (long Bacteria and Archaea, two proteins each).

The reliability filter raises the precision (*Yes* rate) of all four leading methods on short Eukaryotic, Bacterial and Archaeal proteins in every one of the five judge runs (Fig. 4A). These gains are significant in seven of the twelve method-by-group subsets (Fisher exact, *P*_BH_ *<* 0.05). ProtTale’s precision on short

Bacterial proteins rises from 19.2% to 51.2% (*P*_BH_ = 5.7 × 10*^−^*^6^), numerically the largest gain across these two groups.

The same group-specific improvement pattern, after filtering, holds for ProtTale and ProtT3 when predictions are evaluated through GO term extraction using Wang similarity (Fig. 4B) and, less uniformly, other metrics (Extended Data Fig. 8).

### Expert curation validates predictions’ reliability

To check the judge against human expertise, an expert curator, an author not involved in model development, rated ProtTale’s description against the Swiss-Prot description for 232 unseen proteins on a five-point scale, from 1 (unrelated or contradictory) to 5 (equivalent), blind to the reliability score and the judge label (Methods). Expecting mid-range scores to mark text that is hard to judge even for a human, we sampled the tails of the range, *p*_JAY_ ≤ 0.3 and *p*_JAY_ ≥ 0.7, and set aside proteins whose Swiss-Prot description was more than twice as long as the prediction. Thus 90% of what the curator saw (209 of 232) were predictions the model itself did not trust, against 40% of the full unseen set (Supplementary Data 2).

The curator and the judge largely agreed (Extended Data Table 1). The judge said *Yes* to every pair the curator called *equivalent* (5), to 54% of *highly similar* (4) and 22% of *related* (3) pairs, and to only 1% of pairs scored 1–2. Taking a score of ≥4 as agreement, the judge reached 64% precision and 72% recall (86% accuracy; Fisher exact, *P* = 6.3 × 10*^−^*^17^; Spearman *ρ* = 0.585, *P* = 1.0 × 10*^−^*^22^). Its errors sit at the chemically adjacent boundary, with 34% of its *Yes* calls going to pairs scored 3. Of the 56 predictions the curator called *unrelated* (1), the judge labelled 55 not-*Yes*.

**Fig. 3:**
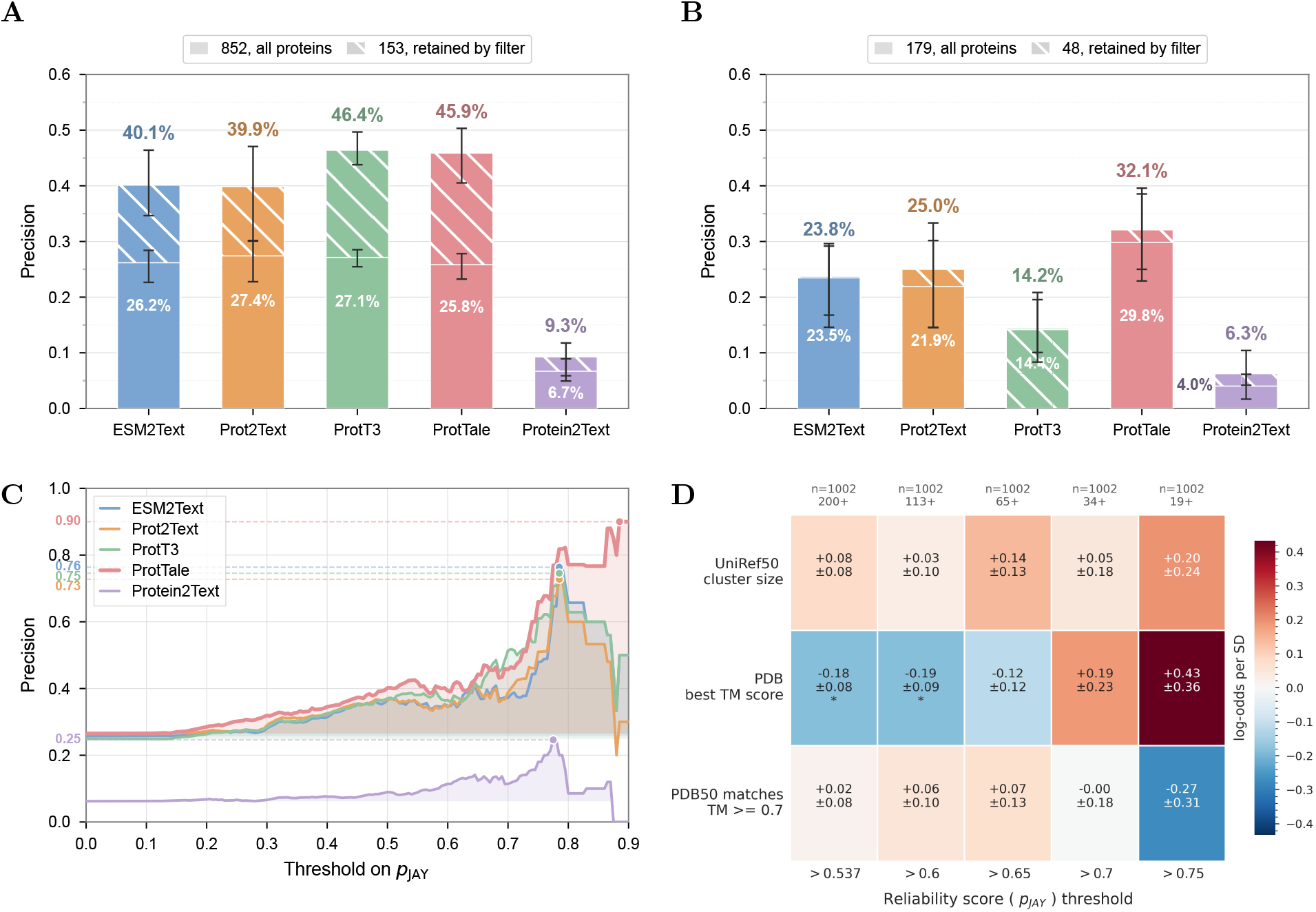
Filtering on the reliability score raises precision for every method on short proteins. *Yes* rate, i.e. precision, with and without the discrete filter *p*_JAY_ *>* 0.537, for **(A)** short (*n* = 852) and **(B)** long (*n* = 179) subsets; *n* is the subset size before filtering; the filter retains the same 153 short and 48 long proteins for every method. **(C)** Continuous filtering, i.e. retaining predictions above a moving *p*_JAY_ threshold rather than a fixed one: precision rises for all four leading methods, but only ProtTale continues to gain at the highest cutoffs, where few predictions are retained. Curves in **(C)** are means over five independent LLM-as-judge runs **(D)** The extent to which known function annotation contributors, i.e. family size and best match in PDB, influence *p_JAY_* reliability score. A logistic-regression (LR) model was trained to predict whether a protein would be retained for ProtTale prediction at each *p_JAY_* threshold. LR derived log-odds scores were computed on the unseen set for: UniRef50 cluster size (sequence family), best TM score vs. PDB100 (structural similarity to experimentally determined structures), and the number of matches in PDB50 with at least TM score of 0.7 (structure family). Mean log odds, with standard error, are reported for each feature, averaged over five LLM-Judge replicates with 10-fold cross-validation. Significance was assessed by Wald tests on the fitted coefficients: *^∗^P <* 0.05, *^∗∗^P <* 0.01, *^∗∗∗^P <* 0.001. Note that regression to predict *p_JAY_* directly failed with poor Mean Absolute Percentage error (≈ 43.6%).

The curator’s scores track predicted reliability (Table 2; Kruskal–Wallis, *P* = 8.0 × 10*^−^*^7^). Above 0.70, more than half the descriptions (57%) were correct and about a tenth (13%) unrelated, so these annotations are worth acting on; below 0.20 the picture inverts, with two thirds (68%) unrelated and a tenth (8%) correct. In every bin the judge’s *Yes* rate stays within five percentage points of the share of predictions the curator scored 4–5, although the head never saw the generated text and the curator never saw the score. Fig. 3C can therefore be read as a guide to what a given score means to a curator.

The most common rating was 3 (82 of 232, 35%), the right functional category with a different specific role. For a *Drosophila* adenylate kinase (Q8IQG9), Swiss-Prot describes reversible phosphate transfer between ATP and AMP, whereas ProtTale predicted phosphorylation of pyrimidine nucleoside monophosphates at the expense of ATP. Both are nucleotide kinases, but the reference acts on a purine (EC 2.7.4.3) and the prediction on pyrimidines (EC 2.7.4.14), which are distinct GO molecular functions, and the judge correctly scored the pair *No*. Such near misses, which are adjacent but still functionally wrong, are what a surface-similarity metric would reward and a reliability filter must catch.

**Fig. 4:**
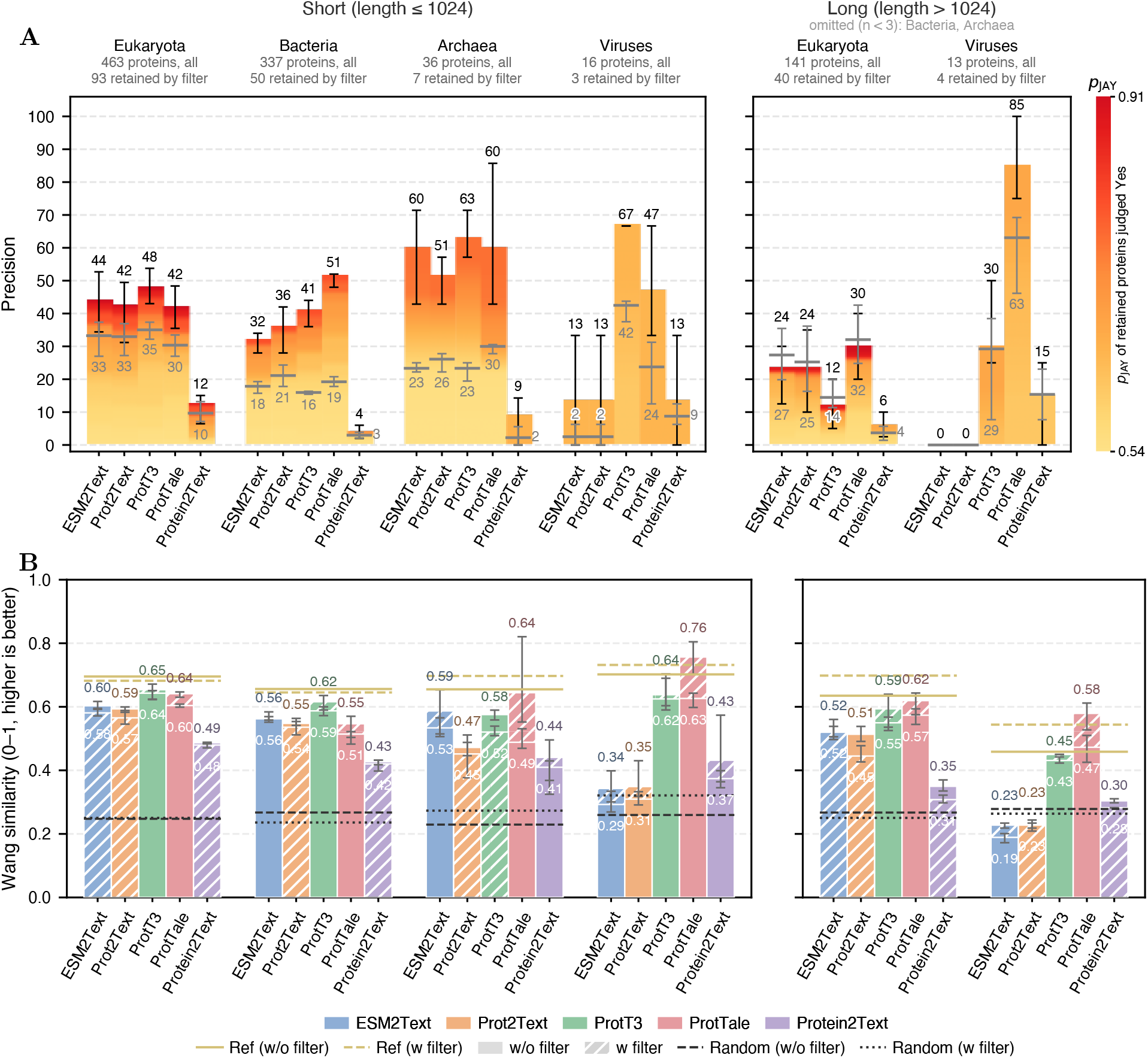
Reliability filtering raises precision in every taxonomic group. Bars and grey lines show the mean across five judge runs, and error bars span the minimum and maximum across those runs. **(A)** Precision with and without the discrete filter (*p*_JAY_ *>* 0.537), by taxonomic group and for short (≤ 1,024 residues) and long (*>* 1,024) subsets. Grey lines give precision on all proteins and bars on the proteins retained by the filter. Bars are coloured by the *p*_JAY_ of the retained proteins judged *Yes*, from 0.54 (yellow) to 0.91 (red), so redder bars draw more of their correct predictions from highly reliable proteins. Long Bacterial and Archaeal subsets retained fewer than three proteins and are not shown. Statistical tests compare filtered with unfiltered results within each method. **(B)** The same comparison on Wang similarity of extracted GO terms (no ancestor expansion), where ProtT3 and ProtTale gain across groups and ESM2Text and Prot2Text gain selectively.

**Table 2:** Expected annotation quality rises with the reliability score. Curator scores for the 232 expert-rated predictions, binned by the head’s output. Note that the curation sampled only the tails of the reliability range (Methods), so no prediction between 0.30 and 0.70 was rated; the gap is a property of the sample rather than of the score. The judge’s *Yes* rate is given beside the curator’s 4–5 rate, which it tracks closely, which is what licenses reading Fig. 3C as a proxy for human quality between the curated tails. Across the full unseen set, 11% of proteins score below 0.20, 28% fall in 0.20–0.30, 57% in the unrated 0.30–0.70 band, and 3% reach 0.70 or above (Supplementary Data 2). Curator scores come from Supplementary Data 1, and reliability scores and judge labels from Supplementary Data 2.

| Reliability score | *n* | Median curator score | Score 1–2 (%) | Score 4–5 (%) | Judge *Yes* (%) |
| --- | --- | --- | --- | --- | --- |
| < 0.20 | 60 | 2 | 68 | 8 | 12 |
| 0.20–0.30 | 149 | 3 | 40 | 19 | 23 |
| ≥ 0.70 | 23 | 4 | 13 | 57 | 52 |

## Discussion

The vast majority of proteins have never been experimentally “seen,” much less functionally annotated. Annotating them by homology can, at best, reproduce what the reference collection already holds and, worse, attach a blatantly wrong label. ProtTale addresses this issue on several fronts. Its supervision does not depend on existing annotations: Machine-Inferred Protein Similarity relies on proximity in ProtT5 embedding space as evidence of shared function, so pair labels can use the whole of the sequence space. ProtTale’s output is not drawn from a fixed vocabulary or pre-determined ontology either, since free text can recombine known functional language instead of picking one catalogued term. Note that this does not escape the problem entirely, as ProtTale learned its language from Swiss-Prot. Together, MIPS-extended supervision and free-text output underlie ProtTale’s distinct coverage of protein space. Skipping the MIPS-supervised alignment that precedes generation halves the share of short proteins whose function the model describes correctly. None of this, however, answers the most salient question a biologist can ask of a prediction: can it be trusted?

Existing protein-to-text models don’t produce a means to tell a well-grounded annotation from a fluent guess. Standard training approaches do not help, as maximizing the likelihood of reference descriptions says nothing about text correctness. We note that a decoder-side signal, e.g. the mean token log-probability of the generated text, is an obvious alternative from the bio-blind world that we did not benchmark. Our reliability head instead scores the protein rather than the text. This parallels pLDDT in structure prediction, where a confidence estimate turns an answer into an answer plus a reason to believe that the model understood the question. This is precisely why our score transfers. Arguably, ProtTale’s biggest function annotation “win” is its ability to say “I don’t know”—and to teach other methods to say it too. Filtering on *p*_JAY_, ProtTale’s reliability score, raises precision for every method we tested on short proteins. What the reliability head reads is therefore not a quirk of ProtTale’s own representation. It is something about the protein, i.e. how readily its mechanism is recoverable from sequence at all.

Our LLM-as-judge protocol scores functional equivalence directly without need for ontologies or token-overlap scores. It assigns *Yes* almost exclusively to functionally matched Swiss-Prot pairs and closely tracks human curator decisions across the decision range. Applied across methods, the judge shows that today’s protein-to-text models reach similar accuracy while succeeding on largely different protein subsets. This orthogonality points to reliability-aware ensembling as the natural next step in annotating functions.

ProtTale has several limitations, none critical to its use, and we aim to address most of them shortly. The judge depends on a particular prompt and evaluator model, and while our five-run analysis addresses run-to-run variation, it cannot rule out drift across model versions; we release all labels and the prompt for re-scoring. Swiss-Prot’s annotation distribution restricts the benchmark to mostly enzymes, leaving functions of metagenome-derived and *de novo* designed sequences mostly untested. Generation caps at 128 tokens, which truncated just under a quarter of the predictions (22.6%), yet the curator rated truncated predictions no lower than complete ones (Mann–Whitney, *P* = 0.25). Running ProtTale over UniProt’s dark and transparent fractions would mean scoring on the order of 10^8^ sequences. At 7,700 sequences per GPU-hour that is about 13,000 GPU-hours, a little over a week on a 64-GPU cluster, and the reliability score comes with it at no extra cost.

As Richard Feynman suggested [53], it is easy to believe findings that match your expectations. A model that writes fluently fools its reader by default, but one that reports its own uncertainty at least makes the deception harder. What our work buys is annotation that an experimentalist can act on: a description for a protein nobody has characterized, and a number saying whether to believe it. More broadly, we propose that pairing generative output with a learned confidence signal is worth asking of any scientific domain.

## Methods

### Machine-Inferred Protein Similarity Score

We embedded 5,190,657 UniRef90 [60] representatives (clusters with ≥10 members, ≤1,200 residues) with ProtT5-XL-UniRef50 [20], mean-pooled and *ℓ*_2_-normalized. Pairs with cosine similarity above 0.77, a threshold at which ≥95% of pairs are real rather than residue-shuffled controls, formed a network of 5.5 billion edges, which we partitioned by Louvain community detection [61] into about 1.2 million clusters. Proteins in the same cluster are labeled functionally similar (positive) and proteins in different clusters dissimilar (negative); these are the MIPS labels used throughout this work. Construction details, and the validation of the clusters against sequence identity, structure and EC numbers, are in Supplementary Note 1.

### Datasets

#### Swiss-Prot source data and splits

We constructed our sequence-to-function set from the full Swiss-Prot database (574,627 entries; release 2026 01), of which 473,996 carry a curated `FUNCTION` description. We took the `FUNCTION` topic of the CC block only, excluding other comment topics such as catalytic activity, subunit, and subcellular location. These were clustered at 40% sequence identity using CD-HIT [40]. All splits below are defined at the level of these clusters, so that no protein in one split shares ≥40% identity with a protein in another. Note that CD-HIT clustering at a 40% identity threshold does not strictly guarantee that every cross-split pair falls below 40% identity; we therefore describe the splits as clustered at 40% identity rather than as an exact identity bound.

##### Unseen evaluation set

All clusters containing proteins deposited into Swiss-Prot after 2022 were assigned to the ***unseen*** set. Because assignment is at the cluster level, a cluster enters the unseen set if *any* member postdates 2022; the set was then restricted to the post-2022 entries themselves, so that every benchmarked protein is itself a post-2022 deposition. No cluster contributed more than one protein, and all other members of these clusters were withheld from training, validation and testing, so no unseen protein shares a 40% identity cluster with any protein the model saw and the benchmark contains one protein per cluster rather than several relatives of the same family. The baseline methods were released at different times and not all of them predate 2022, so a deposition-date cutoff alone does not guarantee that they have not seen these proteins. We therefore removed from the unseen set every protein that appears in the released training data of any baseline evaluated in this work. We note that this removes exact training overlap but, unlike the 40% cluster-level separation applied to ProtTale’s own splits, does not exclude close homologs of baseline training proteins. We further required each protein in the unseen set to have at least one experimentally validated Molecular Function (MF) Gene Ontology (GO) term and at least one Enzyme Commission (EC) number. The EC requirement means the benchmark consists entirely of enzymes, which should be kept in mind when generalizing these results. This procedure yielded 1,031 proteins that serve as a stringent, heldout benchmark. Because ProtTale truncates inputs at 1,024 residues, we report unseen-set results separately for a short subset (≤ 1,024 residues, *n* = 852) and a long subset (*>* 1,024 residues, *n* = 179).

##### Validation and Test sets

After excluding all unseen proteins and their associated clusters, we took the CD-HIT representative of each remaining cluster. Representatives were required to carry at least two experimentally validated MF GO terms and at least one EC number. This is stricter than for the unseen set because these splits train and tune the reliability head, where multi-term proteins give a less ambiguous target; the unseen set stays permissive so as not to favor multi-functional proteins. From this filtered pool, 1,000 were randomly assigned to the validation set and another 1,000 to the test set, and the corresponding clusters were then excluded from further sampling.

##### Training set

From each remaining cluster, up to 10 proteins per cluster were sampled, producing a training set of 207,197 proteins. The cap limits over-representation of large, well-studied families.

#### Stage 1 pairs: functionally similar protein pairs

To pretrain the protein encoder and learn functionally informative embeddings, we constructed pairwise supervision from the machine-inferred protein similarity (MIPS) clusters described in “Machine-Inferred Protein Similarity Score”. Pairs of proteins from the same MIPS cluster were labeled as functionally similar (positive), whereas pairs of proteins from different clusters were labeled as dissimilar (negative). MIPS clusters were built from UniRef90 representatives; each Swiss-Prot protein used here was assigned to the MIPS cluster of its UniRef90 representative, and Swiss-Prot entries with no representative in the MIPS set, i.e. those longer than 1,200 residues or in UniRef90 clusters with fewer than 10 members, contribute no Stage 1 pairs.

Pairs were drawn within the training, validation and test splits defined in “Swiss-Prot source data and splits”, independently in each split to prevent leakage. Within the training split we subsampled 1% of the positive pairs within MIPS clusters, giving 59,773 positive pairs, and drew an equal number of negative pairs from different clusters, giving 119,546 training pairs in total. The same procedure applied to the validation and test splits produced 388 and 534 pairs, respectively (Table 1). These are much smaller because few proteins in a 1,000-protein split share a MIPS cluster. Note that each Stage 1 pair uses both proteins’ curated Swiss-Prot FUNCTION text; it is the *pairing* that is derived without curation, not the text itself.

### Model Architecture

The ProtTale model comprises four components (Extended Data Fig. 1): (1) A protein language model encoder (ESM-C 300M [62]) that maps amino acid sequences to residue-level embeddings, with input sequences truncated to a maximum of 1,024 residues, since attention memory in the encoder grows quadratically with sequence length [63]. (2) A Q-Former projector [64], initialized from PubMedBERT [65] and augmented with cross-attention [63] layers inserted at every transformer block. The projector reads the per-residue embeddings from (1), one vector per amino acid, so that their number changes from protein to protein. Through cross-attention it summarizes this variable-length set into a fixed number of *K* = 4 vectors. (3) A language decoder (Galactica-1.3B [66]) that generates free-text functional descriptions conditioned on the projected query tokens. (4) A reliability classification head that predicts the trustworthiness of each generated description.

### Two-Stage Training Framework

#### Stage 1, Pairwise Protein-Text Functional Alignment

Stage 1 aligns the sequence pathway to a text-derived functional space in a teacher–student set-up (Extended Data Fig. 1A). Each protein of a MIPS-labelled pair contributes its sequence and its curated Swiss-Prot description. A shared Q-Former encodes the two descriptions, and a binary cross-entropy loss on their temperature-scaled cosine similarity, with the MIPS label as target, shapes this text-side space, the teacher. The two sequences, embedded by ESM-C, pass through the same Q-Former, and a pointwise loss (mean squared error plus cosine distance) and a pairwise loss (matching within-batch similarity matrices) pull these sequence-side embeddings, the student, toward the teacher. A stop-gradient keeps the student losses from updating the Q-Former, so the text space is not distorted. The three losses are optimized jointly; their definitions are given in Supplementary Note 5.

#### Stage 2, End-to-End Protein-to-Text Generation

In Stage 2 (Extended Data Fig. 1B), we train an end-to-end model to generate free-text functional descriptions from protein sequences. The model follows an encoder-projector-decoder pipeline conditioned on sequence alone.

Given a protein sequence, we first encode it using the ESM-C protein encoder. The encoder parameters are initialized from the Stage 1 student model and kept frozen during Stage 2 training to preserve Stage 1’s alignment. The resulting residue-level embeddings are passed through the Q-Former, which is trained in this stage, compressing the variable-length protein representation into a fixed number of query output vectors. These query outputs are linearly projected to match the input embedding dimension of the language decoder. The decoder (Galactica-1.3B, fine-tuned with LoRA [67]), together with the Q-Former, is trained to generate Swiss-Prot functional descriptions by maximizing the conditional likelihood of the target text given the protein-derived representations. Specifically, given a target description **y** = (*y*_1_*, . . ., y_T_*) for a protein sequence *x*, we minimize the standard autoregressive cross-entropy loss with teacher forcing (Eq. (1)),

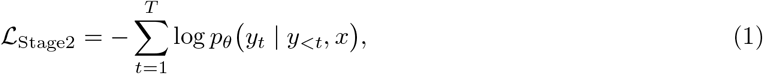

where *p_θ_*denotes the distribution parameterized by the decoder, and *y_<t_* represents preceding tokens. Generation is capped at 128 tokens, both by truncating target descriptions during Stage 2 training and by stopping decoding at inference. We follow ProtT3 [33], whose defaults are the same. The cap affects 26,432 of the 207,197 training targets (12.8%).

The model is trained on the training split and tuned on the validation split (“Swiss-Prot source data and splits”). The test split serves to tune the reliability head (“Reliability Estimation”). All final reporting is performed on the unseen evaluation set.

Inference ran on a single NVIDIA RTX 6000 Ada GPU in bf16 mixed precision with a batch size of 8, annotating the 1,031 unseen proteins in eight minutes, about 7,700 sequences per GPU-hour. This figure includes the reliability head, which adds only a mean-pooling step and one linear layer.

### Existing methods comparisons

We compared ProtTale with four published protein-to-text methods that span the architectural space of the field. ESM2Text, the sequence-only variant released with Prot2Text [38], couples an ESM2 encoder (`esm2_t12_35M_UR50D`) to a GPT-2 decoder and generates from sequence alone. Prot2Text extends the same backbone with a relational graph convolutional network over a residue graph built from the protein’s three-dimensional structure. ProtT3 [33] bridges a frozen ESM2 encoder (`esm2_t30_150M_UR50D`) to Galactica-1.3B through a Q-Former with the query tokens. Protein2Text [39] adapts the LLaVA framework, resampling ESM2-650M residue features into the token space of Llama-3.1-8B-Instruct. All four were run from the checkpoints released by their authors, without retraining or fine-tuning on our data, so each method is evaluated as it is distributed to users.

### GO term extraction from free text

We extracted GO terms from the free-text descriptions generated by each of the five methods, and from the curated Swiss-Prot descriptions used as the curated-text reference, for all proteins of the unseen set. Each description was submitted in a single call to Claude Opus 4.7, rather than through a two-stage extraction-and-grounding pipeline such as OntoGPT [68] (rationale in Supplementary Note 6), prompted to act as a cautious GO curator and to return molecular function identifiers directly supported by the text, or an empty string when the description was too vague to support any (prompt in Supplementary Note 2). Replies were parsed with a regular expression. Extraction was repeated five times per description with independent calls.

For a predicted GO term set *P* and a reference set *R*, term-to-term similarity *s*(·, ·) follows the Wang measure [44] and is aggregated across sets by best-match average,

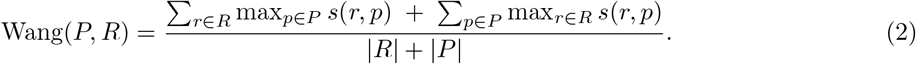

### LLM-as-judge Evaluation

We prompted Claude Opus 4.7, as an LLM judge [47], to act as a Gene Ontology curator and compare each predicted function description with the reference (prompt in Supplementary Note 2). The prompt asked the LLM to focus on molecular function, ignoring cellular localization, biological processes, family membership, or pathway involvement. Each comparison yielded one of four labels reflecting the “curator’s” confidence in functional equivalence,

- *Yes* (score 1) if the predicted vs. reference functions are the same,
- *No* (score 0) if the functions are different,
- *Maybe* (score 0.5) if functions are specifically described but their identity could not be determined,
- *Vague* (score −1) if the prediction was too vague/generic to evaluate.

The *Vague* label penalizes generic outputs such as “binding protein”, and the prompt enforces biochemically meaningful distinctions (e.g., ATP binding ≠ kinase activity) so that superficially similar descriptions are not scored as equivalent. The judge was run five times per prediction, and we considered a prediction to be *Yes* only when at least three of the five runs assigned that label. We used this majority vote for all statistical comparisons of predictions. The 534 Stage 1 test pairs used to validate the judge were scored in a single run.

### Reliability Estimation

Analogous to pLDDT, a reliability head predicts, for each protein, the probability that the judge will label ProtTale’s description *Yes*. Because precision was the goal, *Yes* is the positive class and *No*, *Maybe* and *Vague* are merged into one negative class.

#### Pseudo-labels

After Stage 2 training, the judge scored ProtTale’s predictions on the validation and test splits, and the four-class labels were collapsed to the binary scheme above. Validation labels train the head and test labels tune it, leaving the unseen set for benchmarking (Table 1). The training split is not used, because the generator is likely overfit to it, which would make the labels there heavily imbalanced.

#### Architecture and training

The reliability head takes as input the concatenation of (i) the Q-Former output mean-pooled across query tokens and (ii) the protein encoder output mean-pooled across valid residues. These features are passed through a dropout layer and then through a single linear layer that produces a single logit. Both input features were detached from the computation graph in training, so gradients from the reliability loss did not propagate back to the Q-Former or protein encoder, preserving generation quality. All other model parameters were frozen during reliability head training. Training curves for both stages and the head are in Extended Data Fig. 9 and Supplementary Note 7.

The reliability head was trained using a binary cross-entropy classification loss (Eq. (3)),

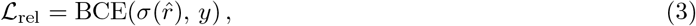

where *r̂* ∈ R denotes the predicted logit, *σ* is the sigmoid function, and *y* ∈ {0, 1} is the binary label with *y* = 1 for *Yes* and *y* = 0 otherwise. At inference, the model returns each description together with *p*_JAY_ = *σ*(*r̂*), the predicted probability of a Judge-Assigned *Yes*, so users can keep only trustworthy outputs without external evaluation. Note that the head is fitted against judge labels, so *p*_JAY_ is a probability by construction; on proteins held out from that fitting it is a prediction.

#### Discrete reliability filter

To anchor where signal begins, we calibrate the head against shuffled amino acid sequences generated by permuting the residues of every protein in the unseen set. A Fisher exact sweep over 401 thresholds on *p*_JAY_ contrasting real and shuffled inputs places the most significant separation at *τ ^∗^* = 0.537, which we adopt as the discrete reliability filter *p*_JAY_ *>* 0.537 throughout the text (procedure in Supplementary Note 4).

#### Evaluation metrics

Precision is the fraction of a set of predictions, all of them or only those the filter retains, that the judge labels *Yes*; filtered predictions do not count against it, because users never see them. Coverage is the fraction of the evaluation set that survives the filter. Sweeping a threshold *t* and retaining predictions with *p*_JAY_ *> t* traces precision against coverage without retraining the generator.

### Expert Curation

One of the authors, a molecular function annotation expert, was recruited to score ProtTale predictions by hand. This gives us a human reference for both the LLM-as-judge and the reliability filter. The expert did not work on the model development nor evaluated the judge prompt.

#### Prediction selection

The predictions rated by the curator were drawn from the unseen set in two steps. First, since a prediction with *p*_JAY_ ≃ 0.5 indicates the head’s lack of confidence, scores in this range cannot be meaningfully evaluated; we therefore restricted the sample to the tails of the reliability range, *p*_JAY_ ≤ 0.3 and *p*_JAY_ ≥ 0.7, which hold 408 and 35 unseen proteins respectively. Second, within those tails we discarded predictions whose Swiss-Prot description was more than twice as long as the prediction, since large length differences could bias the comparison toward differences in annotation detail rather than functional content. This filter left 209 of the 408 and 23 of the 35, giving **232** predictions in total, of which 178 were from the short subset (“Swiss-Prot source data and splits”) with low reliability scores, 18 short with high scores, 31 long with low scores, and 5 long with high scores.

#### Evaluation

Each prediction was presented alongside its Swiss-Prot description in plain text, identified only by Swiss-Prot entry name and sequence length. The reliability score and the LLM-as-judge label were withheld.

Each pair was rated for functional similarity on a five-point scale from 1 (unrelated or contradictory) to 5 (equivalent, the same function and role); the full scale is in Supplementary Note 8.

Ratings reflect what each text states explicitly rather than what could be inferred from it.

### Assessing the effect of sequence representation in function prediction

#### Representation in protein space

To quantify how well each protein in the held-out set is represented in known protein space, we derived three features. First, as a measure of representation in sequence space, we extracted the size (in logarithmic scale) of the UniRef50 (2026 03) cluster[54] containing each protein, which reflects the number of sequence-similar proteins already catalogued. The rest two are measures of representation in structure space. We predicted the structure of each of the 1,002 held-out proteins shorter than 2,750 residues using ESMFold v1 [55] and queried the resulting models against the PDB (250101)[51] at 100% and 50% redundancy (clustered at 50% sequence identity and 80% coverage [56]) using Foldseek [50]. For each protein, we extracted two numbers - the number of matches with at least TM score[57] of 0.7 in PDB50 and the best TM-score (alntmscore) [50] in PDB100, as indicators of whether a structurally similar protein is already known. Together, these three features serve as proxies for homology to previously characterized proteins, and we used them to test the influence of prior representation on function prediction across models and on ProtTale’s reliability score.

#### Feature-contribution models

We fitted a logistic regression relating the three representation features to the verdict of an LLM judge, separately for each method and judge run. The three features were standardized to zero mean and unit variance. Because the judge scored every description five times, each fit was assessed by its own stratified 10-fold cross-validation [58], yeilding fifty held-out estimates per method. The reported coefficient is the mean across the five runs; its standard error combines the within-run model-based error with the spread of the five estimates, added in quadrature. The significance was assessed using Wald tests on the average coefficients [59], and each coefficient is expressed as the change in log-odds of a correct description per standard deviation of the feature. We also repeated the experiment with reliability score-derived judgement in place of the LLM judge, to assess the influence of homology on the reliability module.

### Statistical analysis

All tests are two-sided at *α* = 0.05, with *P* -values corrected within each family of related tests. Because every method predicts every unseen protein, labels from different methods are paired, and method comparisons use paired tests.

For the GO-extraction comparison (Results, “Mapping free text to GO terms”), we averaged per-protein Wang similarity over the five extraction runs, scoring empty extractions as zero so that proteins stay paired, and compared all ten method pairs with Wilcoxon signed-rank tests and Holm–Bonferroni correction. Judge accuracy was tested with Fisher’s exact test on the 534 Stage 1 test pairs, same versus different MIPS cluster. Judge reproducibility was summarized by agreement and Cohen’s *κ* over the ten pairs of runs per method, with 95% bias-corrected and accelerated [69] bootstrap intervals from 10,000 resamples.

Method pairs were compared with McNemar’s test [70], using the continuity-corrected *χ*^2^ approximation when the two methods disagreed on at least 25 proteins and the exact binomial test otherwise, for all ten pairs before filtering (Results, “Free-text predictors”) and, after filtering, for one pair on short and four pairs on long sequences (Results, “The reliability head scores the protein”). The effect of filtering within a method was tested with Fisher’s exact test comparing retained with rejected predictions, so that the two groups are disjoint, with Holm–Bonferroni correction over the ten short and long tests and Benjamini–Hochberg correction over the thirty group-by-length tests (Results, “Reliability filtering helps in every taxonomic group”). Figures instead plot the retained subset against the full subset.

For expert curation, a Mann–Whitney *U* test compared curator ratings of truncated and complete predictions, counting a prediction as truncated when generation reached the 128-token cap without an end-of-sequence token. A Kruskal–Wallis test compared ratings across the reliability bins of Table 2, and Spearman’s *ρ* measured their rank correlation with judge labels. Conclusions are unchanged after correction for multiple testing.

## Supplementary information

This article contains Extended Data Figs. 1–9, Extended Data Table 1, Supplementary Notes 1–8 and Supplementary Data 1–7, whose contents are listed under Supplementary Data.

## Acknowledgements

This work was supported by NASA Astrobiology Institute grant 80NSSC18M0093 and by the National Science Foundation (NSF) awards 1553289 (CAREER) and 2310114 to Y.B.

## Declarations

### Code availability

ProtTale is available as a web service at https://services.bromberglab.org/prottale/. Source code for training, inference, and all analyses reported here is available at https://github.com/mulahteele/ProtTale under an MIT license. The trained model checkpoint is available at https://huggingface.co/Mulah/ProtTale/blob/main/checkpoint.ckpt under a CC BY-NC 4.0 license, inherited from its Galactica-1.3B backbone.

## Data availability

The Stage 1 and Stage 2 Swiss-Prot splits, the 40% CD-HIT clustering that defines them, and Supplementary Data 1–7 are available at https://huggingface.co/Mulah/ProtTale/tree/main/data under a CC BY-NC 4.0 license. Swiss-Prot annotations were obtained from UniProtKB (https://www.uniprot.org).

## Author contributions

T.L. implemented the model, ran all experiments, processed and analysed the data, and wrote the first draft of the manuscript. R.P. advised on the direction of the experiments and contributed to the writing. R.T. performed the blinded expert curation of model predictions. F.G. evaluated and validated the machine-inferred protein similarity clustering. C.Y. advised on the model architecture.

Y.B. conceived the study, supervised the work throughout and revised the manuscript. All authors read and approved the final manuscript.

## Extended Data

**Extended Data Fig. 1:**
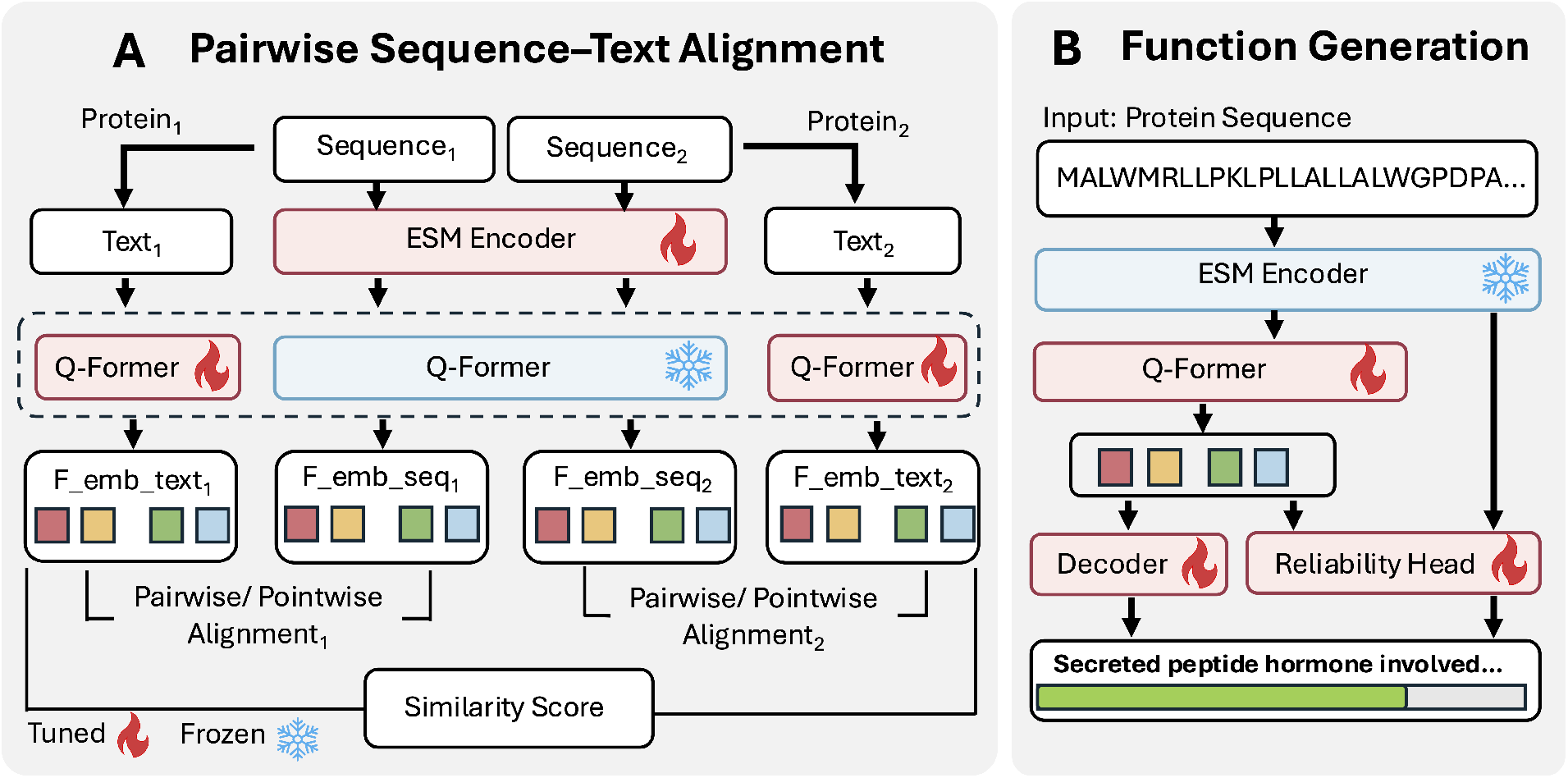
Overview of the ProtTale architecture. Flames mark tuned components, snowflakes frozen ones. **(A)** Stage 1, pairwise sequence–text alignment. Each protein of a MIPS-labelled pair contributes its amino acid sequence, embedded by the ESM encoder, and its curated Swiss-Prot description. A shared Q-Former encodes both sides, tuned on the text pathway and frozen on the sequence pathway, so that sequence embeddings are pulled toward the text-derived space by pointwise and pairwise alignment losses against the pair’s similarity label. **(B)** Stage 2, function generation from sequence alone. The ESM encoder carries over the weights learned in Stage 1 and is frozen, while the Q-Former and decoder are tuned to generate free text. A reliability head reads the encoder output and estimates how likely the LLM-as-judge would be to call the generated description a match (green bar).

**Extended Data Fig. 2:**
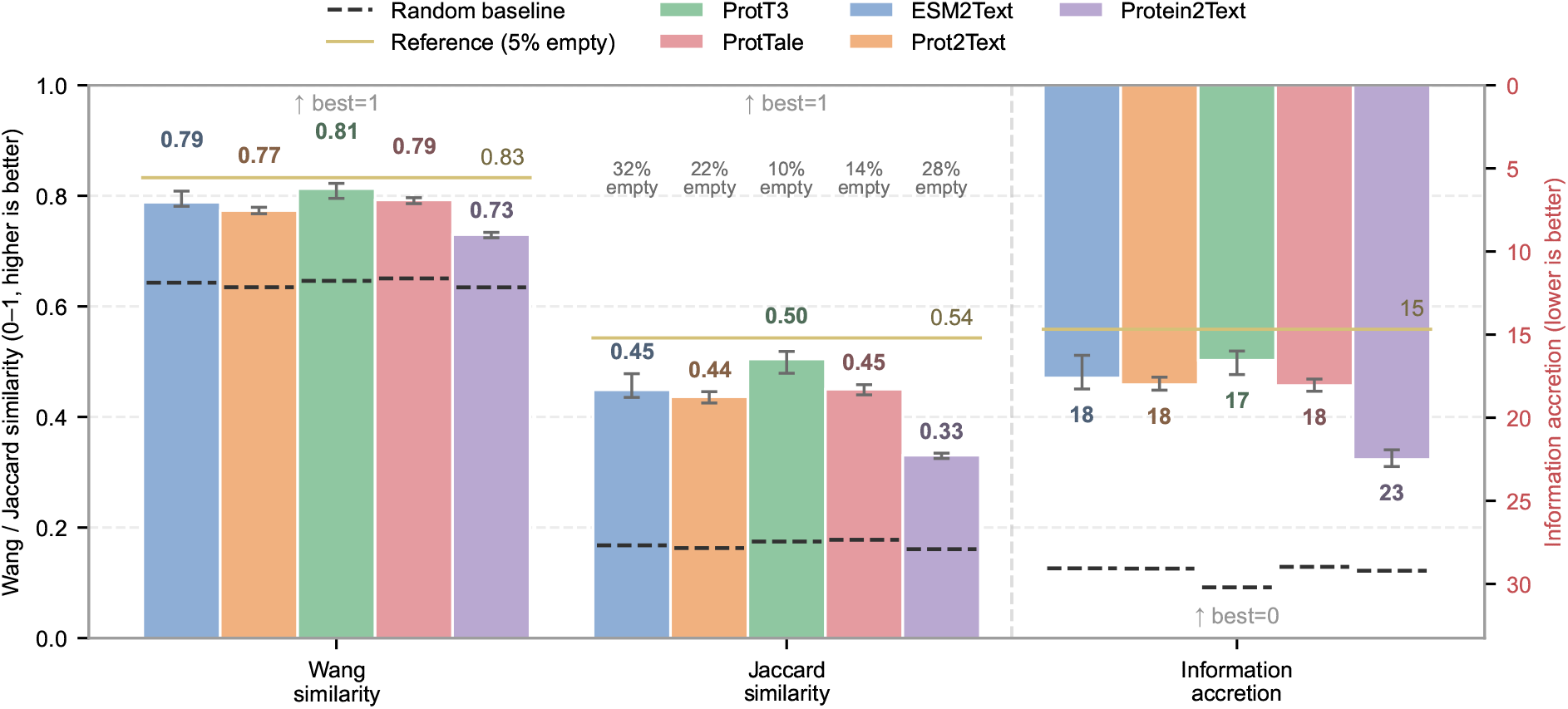
Ancestor expansion does not change the picture: methods remain clustered below the curated-text reference. Wang similarity, Jaccard similarity, and information accretion on the unseen evaluation set after ancestor expansion of GO terms. Bars are per-method means over five runs, with error bars spanning the minimum and maximum across runs. The solid olive line marks the curated-text reference from the curated Swiss-Prot description, the dashed black line marks the random baseline, and percentages above bars are empty-extraction rates.

**Extended Data Fig. 3:**
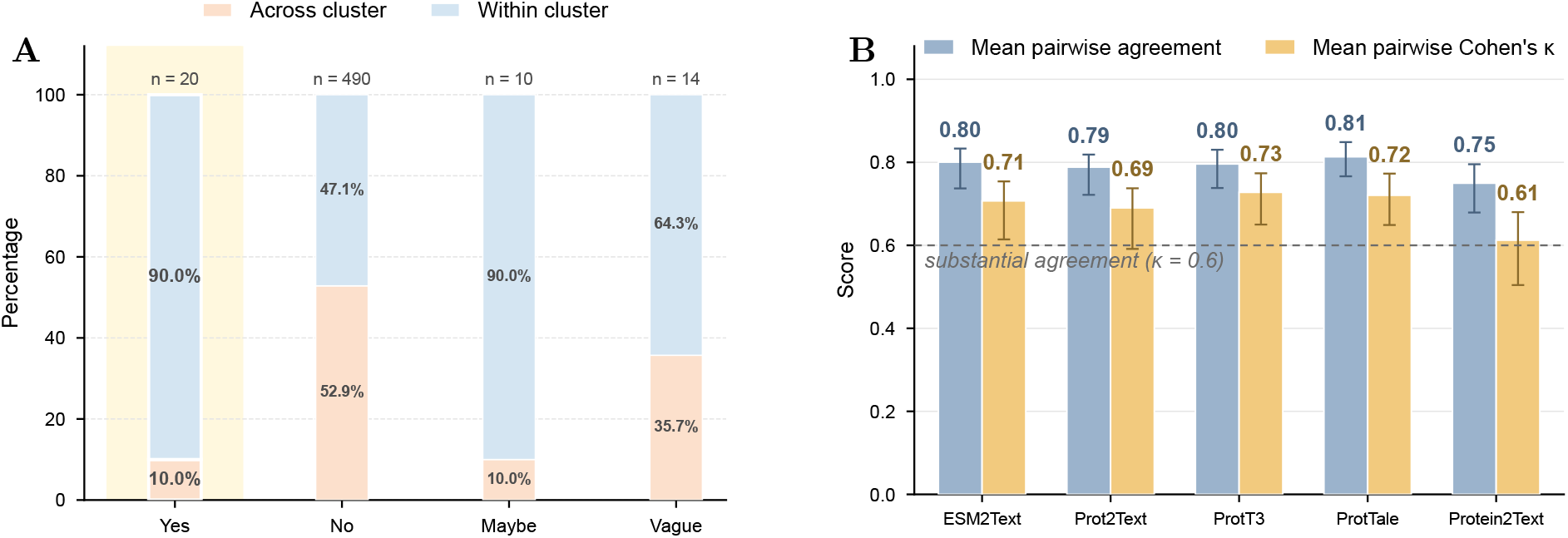
LLM-as-judge is biologically precise, if conservative, and reproducible across runs. (**A**) Judge labels, from a single run, for the 534 Stage 1 test-set protein pairs: 267 within-cluster pairs, expected to share function (blue), and 267 across-cluster pairs (orange). Each label column is normalized to 100%, so the split within a column gives the fraction of pairs with that label from the same cluster; counts appear above each bar. (**B**) Agreement between five independent judge runs on the unseen-set predictions, as Cohen’s *κ*. Error bars span the 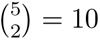 run pairs, and the dashed line marks the “substantial” threshold *κ* = 0.6.

**Extended Data Fig. 4:**
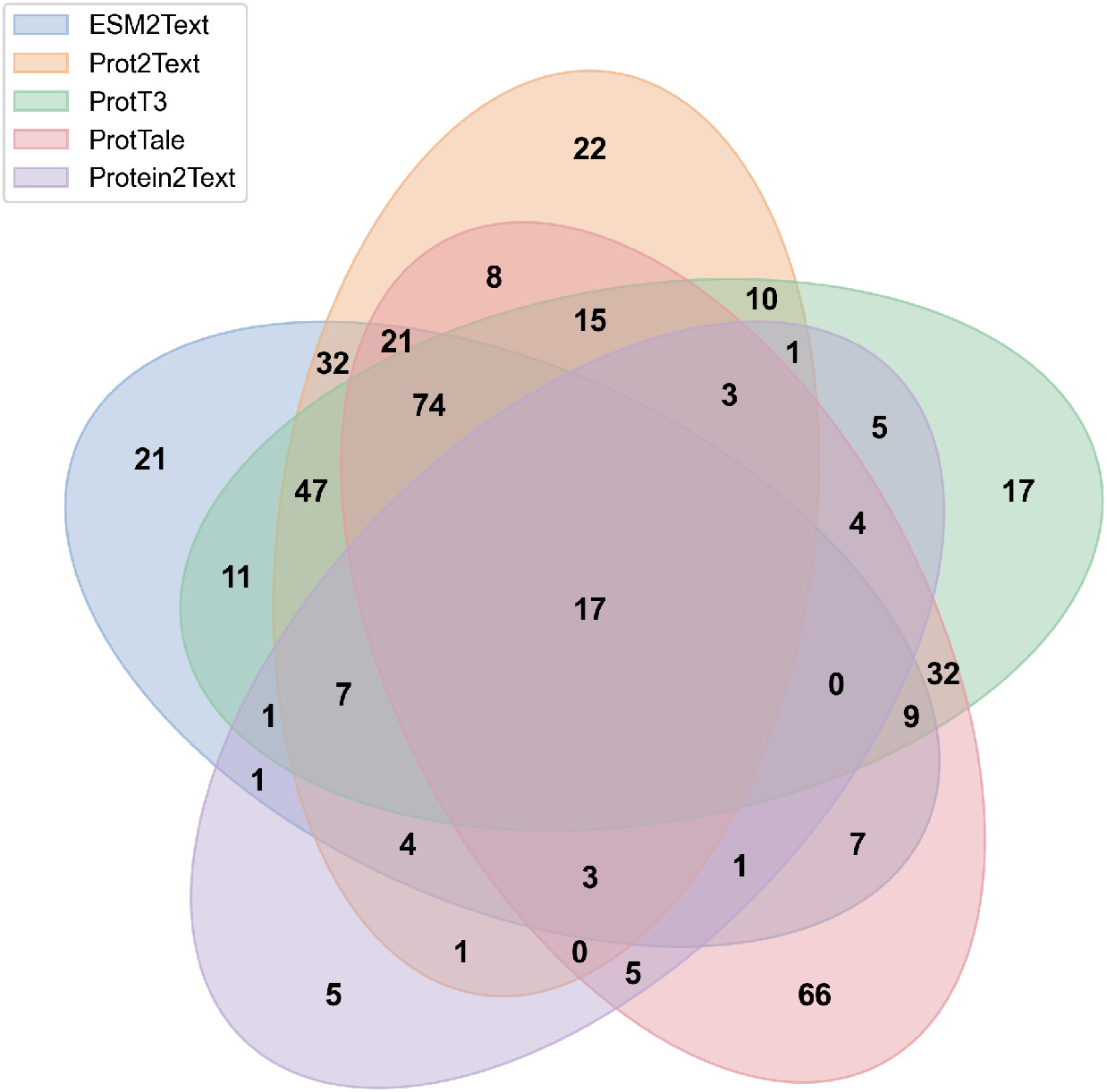
Overlap of *Yes* predictions across all five methods. Five-way Venn of proteins labelled *Yes* on the unseen evaluation set, adding ESM2Text and Protein2Text to the three-method comparison of Fig. 2B under the same judge criterion. The central region (17) contains proteins recovered by all five methods. Single-method slices (21, 22, 17, 66 and 5 for ESM2Text, Prot2Text, ProtT3, ProtTale and Protein2Text) contain proteins recovered by exactly one method. ProtTale’s unique set is smaller here than in the three-method comparison of Fig. 2B, since adding two further methods claims some of those proteins.

**Extended Data Fig. 5:**
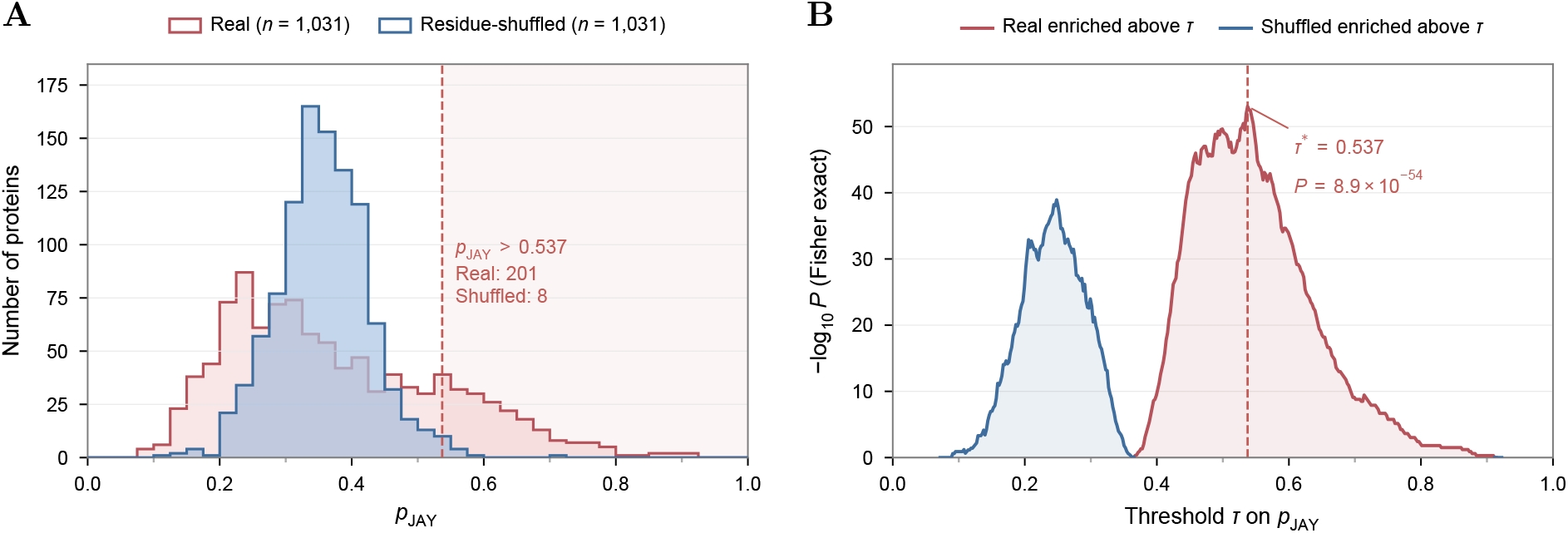
Real proteins separate from residue-shuffled controls, fixing the discrete filter at τ ∗ = 0.537. Shuffled amino acid baseline for the ProtTale reliability head on the unseen evaluation set. **(A)** *p*_JAY_ distributions for the real unseen sequences and their residue-shuffled versions (*n* = 1,031 each). The shaded region marks the discrete filter *p*_JAY_ *>* 0.537, which retains 201 real and 8 shuffled proteins. **(B)** Fisher exact test of real against shuffled proteins at each threshold *τ* on *p*_JAY_, in red where real proteins are enriched above *τ* and in blue where shuffled proteins are. Significance peaks at *τ ^∗^* = 0.537 (dashed line), which we adopt as the discrete reliability filter in the main text.

**Extended Data Fig. 6:**
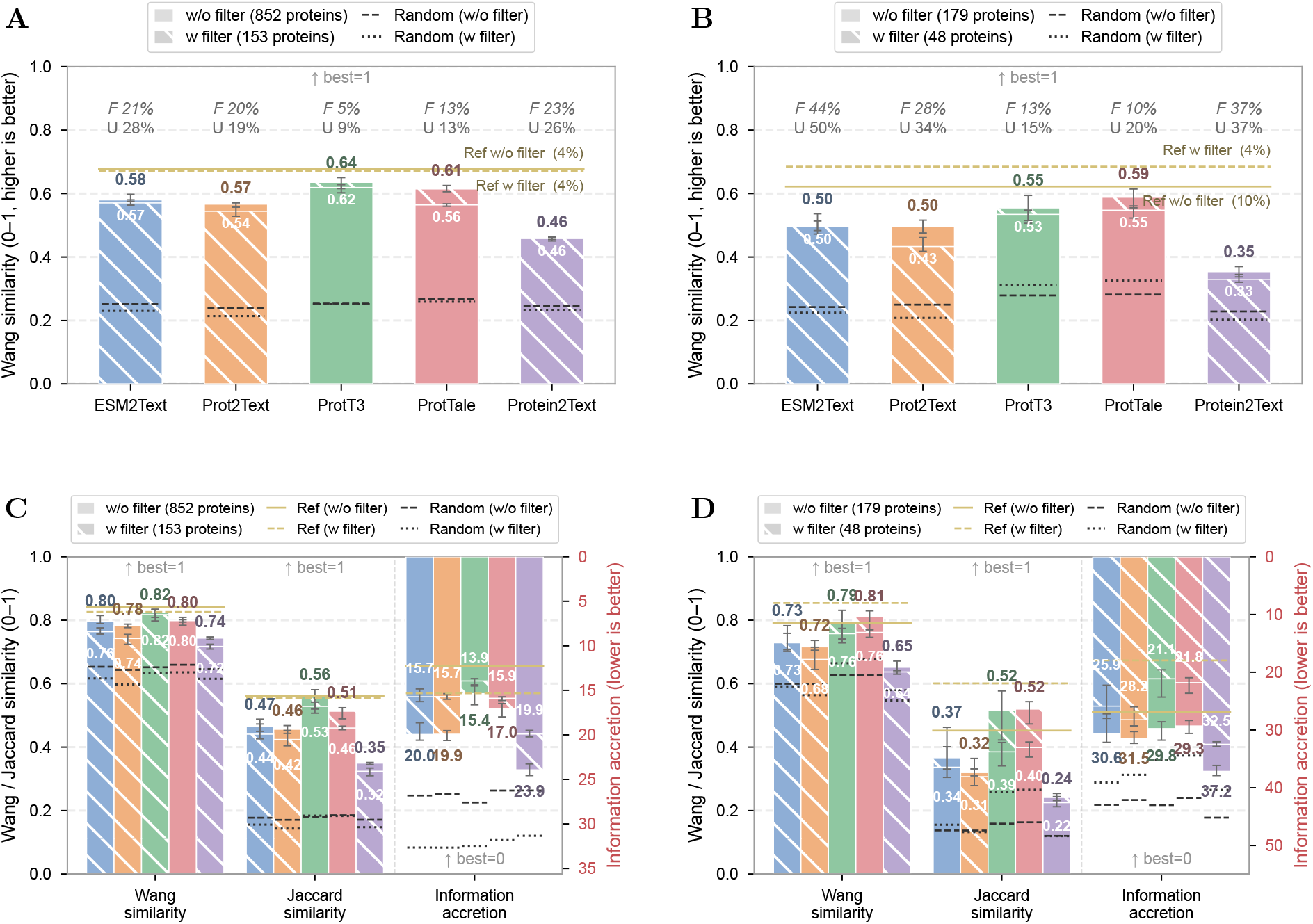
The reliability filter improves Wang similarity with and without ancestor expansion. Wang similarity of extracted GO terms with and without the discrete filter *p*_JAY_ *>* 0.537, for the short (*n* = 852) and long (*n* = 179) subsets of the unseen evaluation set. **(A,B)** Without ancestor expansion, as reported in the main text; only ProtT3 and ProtTale gain. **(C,D)** With every GO term expanded to its ancestors before scoring, which mirrors the no-expansion result, with ProtTale’s lead largest on the long subset.

**Extended Data Fig. 7:**
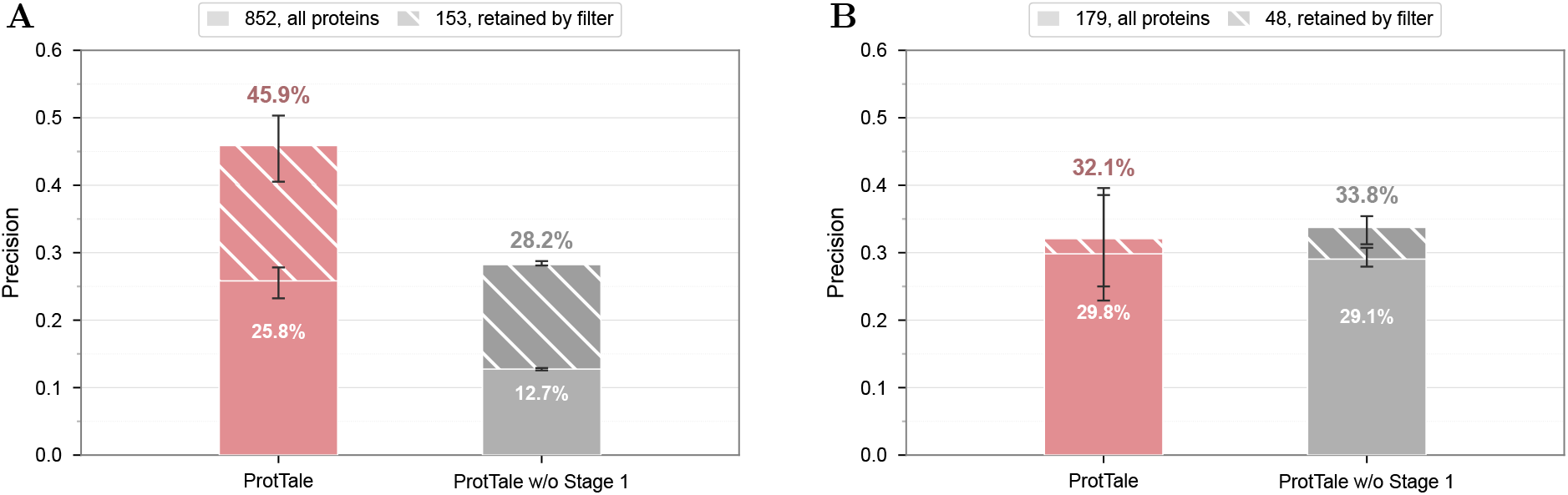
Removing Stage 1 halves precision on short proteins and leaves long proteins unchanged. Precision of ProtTale and of ProtTale retrained without Stage 1 (ProtTale w/o Stage 1) on the **(A)** short (*n* = 852) and **(B)** long (*n* = 179) subsets of the unseen set, with and without the discrete filter *p*_JAY_ *>* 0.537, computed by ProtTale’s reliability head for both models. ProtTale w/o Stage 1 was trained with Stage 2 only, its frozen encoder initialized from the original ESM-C weights rather than from the Stage 1 student, with every other setting unchanged, and was scored with the same five-run judge protocol. Bars are means over five judge runs and error bars span the minimum and maximum across runs.

**Extended Data Fig. 8:**
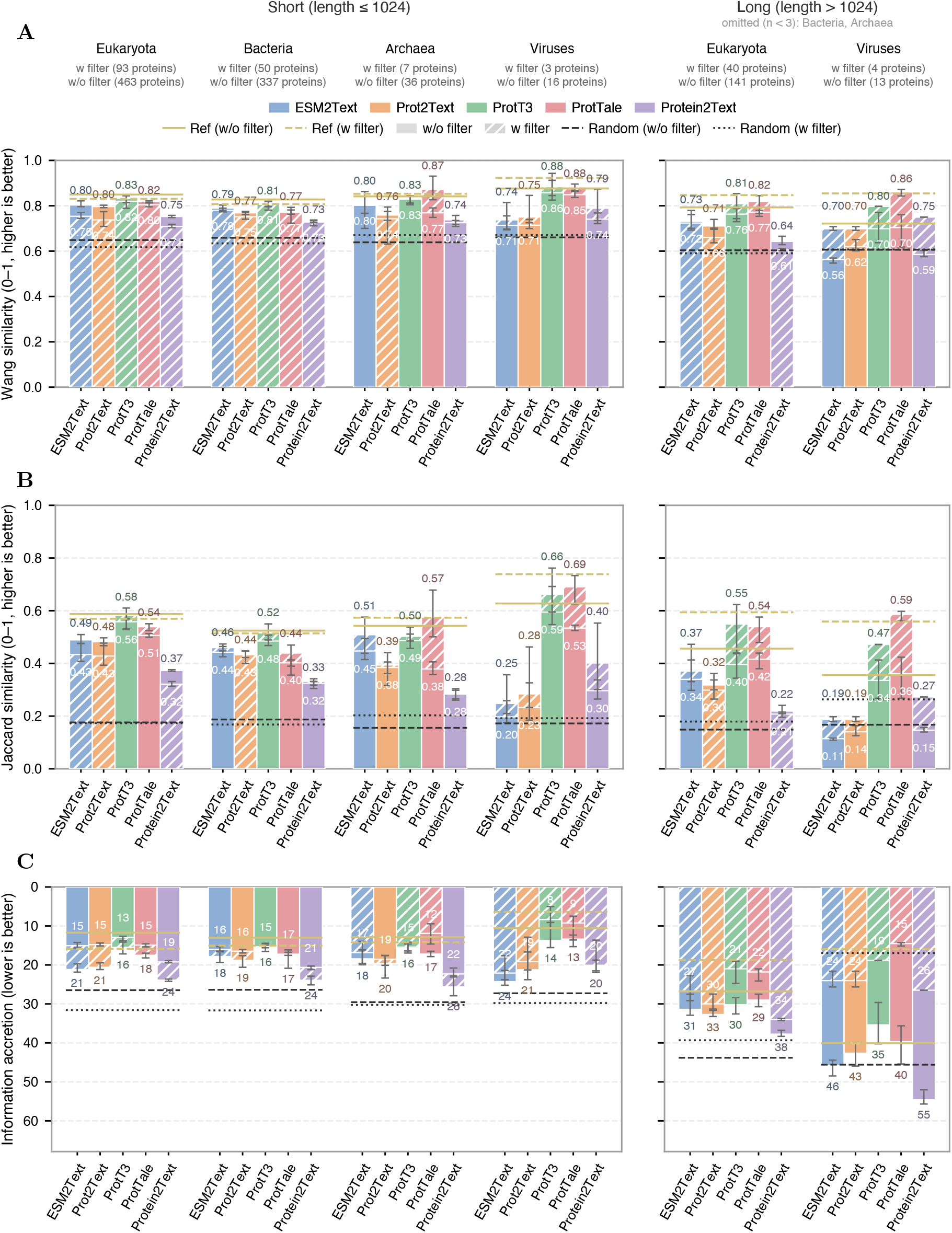
With ancestor expansion, per-group filter gains are less uniform. The counterpart to Fig. 4B, with every GO term expanded to its ancestors before scoring. ProtTale and ProtT3 improve Jaccard similarity in every retained group and Wang similarity and information accretion in four of six, while the other methods gain in fewer groups. Per-group analysis under *p*_JAY_ *>* 0.537. **(A)** Wang similarity, **(B)** Jaccard similarity, and **(C)** information accretion with and without the filter per method, on short and long subsets of the unseen evaluation set.

**Extended Data Fig. 9:**
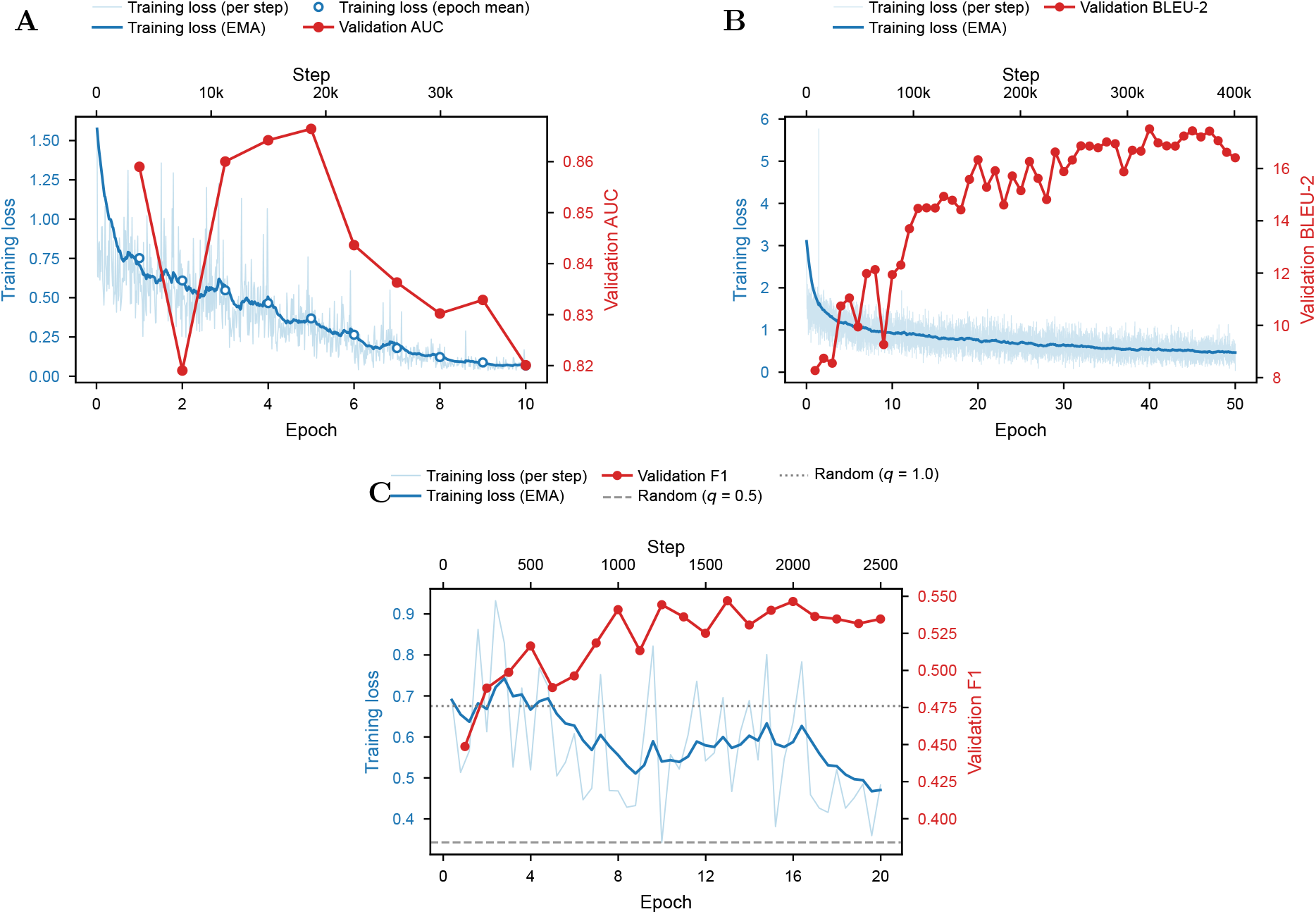
Training dynamics for both stages and the reliability head. Curves for the two ProtTale training stages and the reliability head. Each panel shows training loss (left, blue) and the validation metric (right, red). **(A)** Stage 1 alignment, validation sequence AUC across 10 epochs. **(B)** Stage 2 generation, validation BLEU-2 across 50 epochs. **(C)** Reliability-head fine-tuning, validation positive-class F1 across 20 epochs with random baselines at *q* = 0.5 (0.384) and *q* = 1.0 (0.476), where *q* is the rate at which the random classifier predicts the positive class.

**Extended Data Table 1:** The LLM-as-judge agrees with a blinded human curator at the extremes. Fraction of the 232 expert-rated predictions that the judge labelled *Yes*, by the curator’s 1–5 similarity score.

| Curator score | $n$ | Judge called it *Yes* |
| --- | --- | --- |
| 1 — unrelated or contradictory | 56 | 2% |
| 2 — somewhat related | 47 | 0% |
| 3 — related, distinct role | 82 | 22% |
| 4 — highly similar | 28 | 54% |
| 5 — equivalent | 19 | 100% |

## Supplementary Information

### Supplementary Note 1: Construction and validation of the MIPS network

#### Sequence and embedding set

We compiled a dataset of 5,190,657 representative protein sequences from UniRef90 [60] clusters with at least 10 members and a maximum sequence length of 1,200 amino acids. Per-protein representations were generated using ProtT5-XL-UniRef50 [20], a pre-trained protein language model that captures evolutionary, structural, and functional constraints. Each protein sequence was mapped to a 1,024-dimensional vector by averaging the residue-level ProtT5 embeddings, which was then *ℓ*_2_-normalized to unit length.

#### Network of functionally similar protein pairs

We constructed a protein similarity network where nodes represent protein sequences and edge weights correspond to cosine similarity between their embeddings. Computing the full O(*N* ^2^) pairwise similarity matrix for over 5 million sequences (∼13 trillion edges) is computationally infeasible, so we retained only edges likely to capture functional relevance. To set the pruning threshold, for a subset of 1,000 proteins we compared cosine similarity distributions between *Real vs. Real* protein pairs and *Random vs. Real* pairs, the latter generated by randomly shuffling the amino acid sequence of a corresponding real protein. A cosine similarity threshold of 0.77 was selected such that ≥95% of pairs exceeding this value came from the Real– Real distribution. Applying this threshold with FAISS FlatIP vector search [71] gave a sparse network of 5,540,906,042 edges (∼0.04% of the fully connected set).

#### Clustering

To identify functionally coherent protein groups, we used a multi-stage clustering pipeline implemented with RAPIDS cuGraph. A random subsampling strategy applied Louvain community detection [61] iteratively to manageable subgraphs while ensuring full node coverage. Cluster assignments from multiple iterations were consolidated by a Jaccard similarity-based consensus alignment, and a refinement procedure then split overly large clusters to improve functional specificity. The code is in the ProtTale repository.

#### Validation

To verify that the clusters reflect biological similarity, we sampled 1,000 same-cluster protein pairs, restricted to Swiss-Prot entries with an assigned EC number and an available AlphaFold structure, capping selection at five pairs per cluster so that large clusters do not dominate (244 clusters represented). For each pair we computed pairwise sequence identity by global alignment (BLOSUM62; gap open −10, extend −0.5, normalized by the shorter sequence length), structural similarity by TM-align [49] on AlphaFold Protein Structure Database structures [73], and EC number concordance at the fourth, fully specified level. Per-pair values and summary statistics are in Supplementary Data 5.

### Supplementary Note 2: Prompts

#### GO term extraction

The ontology-based baseline analysed in Results, “Mapping free text to GO terms”, extracts GO terms from a free-text protein description with the prompt below. It directs the model to act as a cautious GO curator that prefers high precision over recall, and to return only GO identifiers directly supported by the input description. The user message inserts a single description per call into the `{sentence}` placeholder.

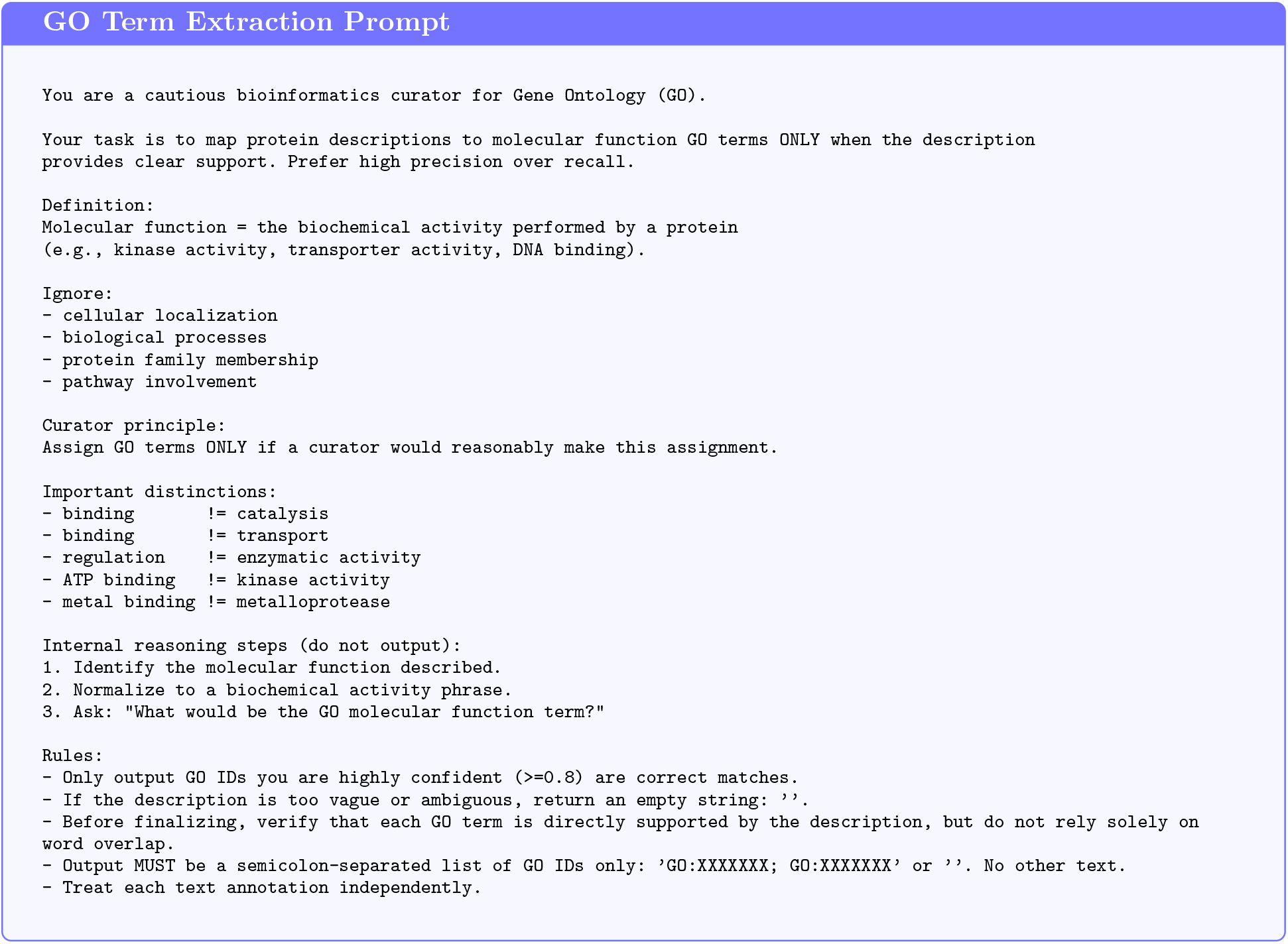

#### LLM-as-judge

The judge compares a predicted protein description with its reference and returns one of four labels (Methods, “LLM-as-judge Evaluation”). The prompt directs the model to act as a Gene Ontology curator focused exclusively on molecular function, to ignore other ontology branches such as cellular localization or biological process, and to output a numerical score mapping to *Yes*, *No*, *Maybe* or *Vague*.

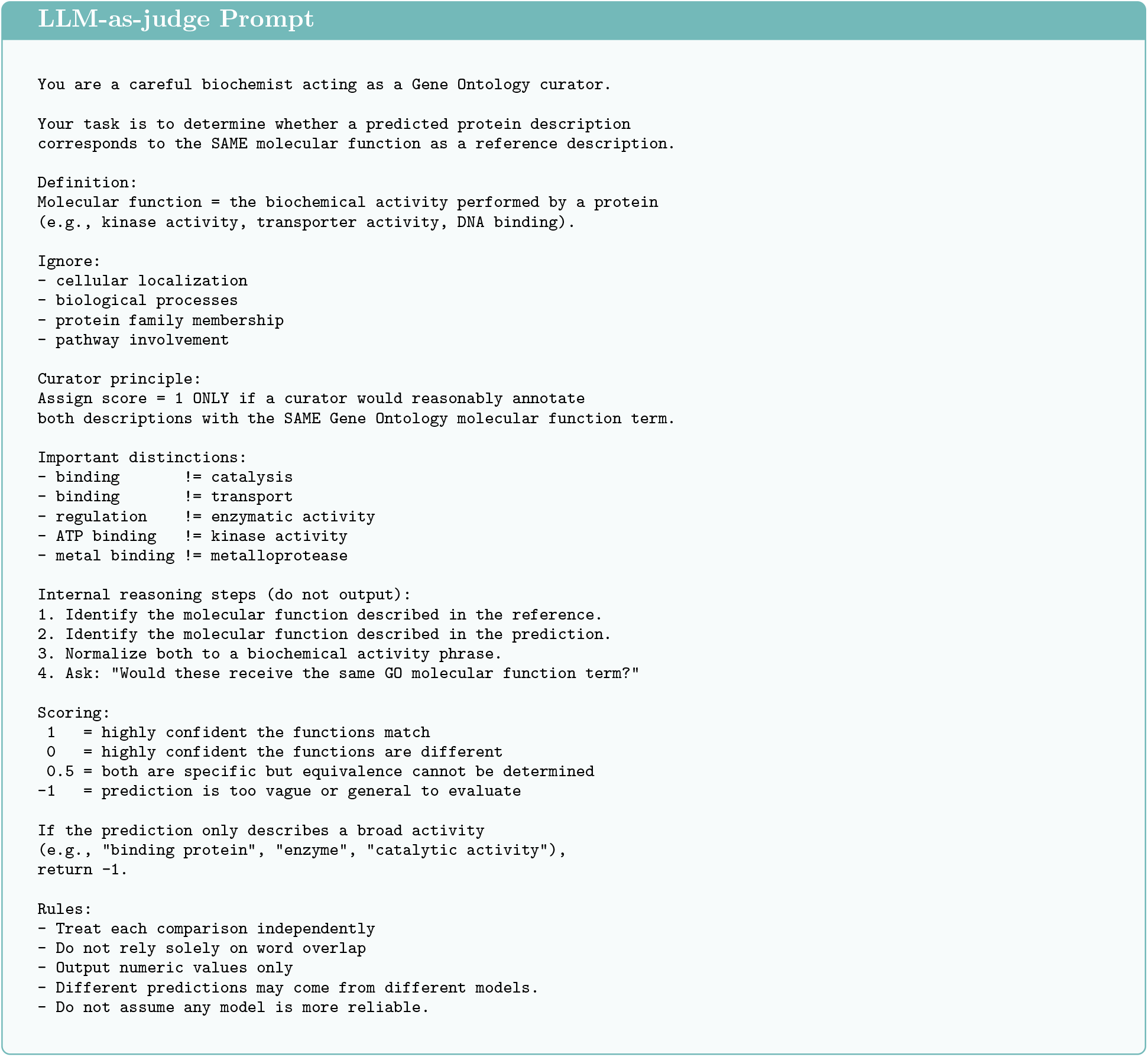

### Supplementary Note 3: GO term extraction with ancestor expansion

Results, “Mapping free text to GO terms” of the main text reports Wang semantic similarity without ancestor expansion. We additionally evaluated the same extracted GO sets after expanding every term to its ancestors and rescoring with three complementary metrics, Wang similarity, Jaccard similarity, and information accretion (Extended Data Fig. 2).

The expanded view is consistent with the no-expansion view in the main text. All four leading methods cluster within a narrow window across all three metrics, with Wang similarity from 0.77 to 0.81, Jaccard similarity from 0.44 to 0.50, and information accretion from 16.5 to 18.1. ProtT3 leads marginally on Wang and Jaccard while ESM2Text and ProtTale tie within error bars, and information accretion separates Protein2Text from the rest but leaves the four leading methods close together. The curated-text reference at 0.83 Wang, 0.54 Jaccard, and 14.7 information accretion remains close to the leading methods, so ancestor expansion does not enlarge the gap that ontology mapping introduces, leaving the methods indistinguishable.

### Supplementary Note 4: Shuffled amino acid baseline for the reliability head

To verify that the reliability head separates real proteins from shuffled inputs, we generated a negative set by randomly permuting the residues of every unseen protein and ran ProtTale on both real and shuffled inputs. The two *p*_JAY_ distributions diverge in the upper tail (Extended Data Fig. 5A). Shuffled sequences concentrate around *p*_JAY_ ≈ 0.35, and only 8 of 1,031 exceed 0.537, while real proteins extend a long right tail past *p*_JAY_ = 0.90. Sweeping a threshold *τ* on *p*_JAY_ and computing a Fisher exact test that contrasts the proportion of inputs above *τ* between the two populations, the most significant separation is reached at *τ ^∗^* = 0.537 (*P* = 8.9 × 10*^−^*^54^, Extended Data Fig. 5B). We adopt *τ ^∗^* = 0.537 in the main text (Results, “The reliability head scores the protein”) as the discrete reliability filter, so any retained prediction must score above what shuffled noise would receive. At each of 401 evenly spaced thresholds *τ* we tabulated real and shuffled sequences into a 2 × 2 table of (real, shuffled) × (above *τ*, at or below *τ*) and applied a two-sided Fisher exact test, taking *τ ^∗^* as the smallest *p*-value among thresholds where more real than shuffled sequences fell above *τ* . As the minimum over a sweep, this *p*-value is a descriptive optimum rather than a corrected significance level.

### Supplementary Note 5: Stage 1 alignment objectives

For the text pathway, each functional description in a pair is encoded by the Q-Former, with weights shared between the two members of the pair (a siamese configuration), producing text-side functional embeddings. The Q-Former is optimized with a binary cross-entropy loss L_BCE_ that uses the temperature-scaled cosine similarity between the two text-side embeddings as its logit and the MIPS label as its target, so same-cluster descriptions are driven together and different-cluster descriptions apart. This text-derived space serves as the *teacher*.

For the sequence pathway, the sequences of the same pair are encoded by the ESM-C protein encoder, again with shared weights, and passed through the *same* Q-Former to obtain sequence-side embeddings, the *student*. Gradients from the two alignment losses are stopped at the Q-Former on this path, so the sequence pathway projects into the text-derived space without distorting it; the Q-Former itself is updated only by the text-side objective.

The pointwise alignment loss enforces consistency between sequence-side and text-side embeddings through a weighted combination of mean squared error and cosine distance,

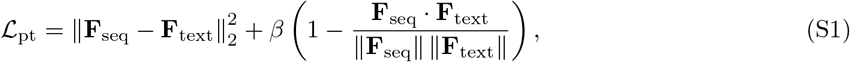

where *β* controls the weight of the cosine term. The pairwise alignment loss preserves relative functional relationships within a mini-batch by matching pairwise cosine similarity matrices,

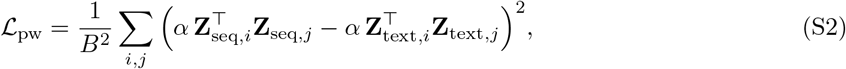

where **Z**_seq_ and **Z**_text_ are *ℓ*_2_-normalized sequence and text embeddings and *α* is a scale applied to both similarity matrices. Because *α* factors out of the squared difference as *α*^2^, it is not separately identifiable from *λ*_pw_ in Eq. (S3); we fix *α* and tune *λ*_pw_ only. All three terms are optimized jointly in a single objective; the teacher–student asymmetry comes from the stop-gradient, not from an ordering of training runs:

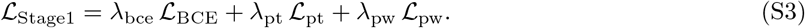

### Supplementary Note 6: Choice of a single-step GO extraction

Ontology-grounded extraction frameworks such as OntoGPT [68] provide a natural approach to mapping text to GO terms, first extracting ontology-relevant entities with an LLM and then grounding them to standardized GO identifiers through ontology annotators. Such pipelines were developed for literature-scale narrative text, whereas Swiss-Prot function descriptions are short and densely packed, often stating several overlapping activities in a single sentence, so entity boundaries are poorly defined and errors at the extraction stage carry into grounding. We therefore bypassed the two-stage pipeline and prompted a single model to assign GO terms to a description directly.

### Supplementary Note 7: Training dynamics

We report training curves for the two ProtTale training stages and, separately, for the reliability head, which is fitted on frozen features after generative training is complete (Extended Data Fig. 9). Every panel shows training loss on the left axis and the primary validation metric on the right axis.

#### Stage 1, pairwise protein to text alignment

In Stage 1 the ESM-C encoder is trained to map sequence-derived embeddings into the text-derived functional space under pairwise MIPS supervision, with the Q-Former acting as a fixed projection on this path (Methods, “Stage 1, Pairwise Protein-Text Functional Alignment”). Validation sequence AUC peaks at approximately 0.87 around epoch 5.

#### Stage 2, end-to-end protein to text generation

With the Stage 1 encoder frozen, the Q-Former and the LoRA adapters on the Galactica-1.3B decoder are optimized for Function description generation. Validation BLEU-2 peaks at approximately 17 around epoch 40.

#### Reliability head fine-tuning

All other model parameters are frozen and the head’s input features are detached, so the head cannot interfere with generation quality. Positive-class F1, measured on the test split (which serves as the reliability head’s tuning set; Table 1) while the head is fitted on the validation split, peaks at approximately 0.55 around epoch 13, above both random baselines, where *q* denotes the probability with which a random classifier predicts the positive class (*q* = 0.5 gives F1 = 0.384; *q* = 1.0, i.e. always predicting positive, gives F1 = 0.476, implying a positive-class prevalence of about 31%), and we adopt this checkpoint for all reliability results in the main text (Extended Data Fig. 9C).

### Supplementary Note 8: Expert curation of ProtTale predictions

The complete per-prediction curation is provided as Supplementary Data 1 (`Supplementary_Data_1_expert_curation.csv`), with one row for each of the 232 curated predictions. Sampling and blinding are described in Methods, “Expert Curation”.

#### Protein

Entry is the identifier shown to the curator, running from 1 to 232 in the order the predictions were presented. `UniProt_ID` is the Swiss-Prot entry name and `Length_aa` the sequence length.

#### Model output

`p_JAY` is the reliability score assigned by ProtTale’s head, and `Retained` records whether it passes the discrete filter *p*_JAY_ *>* 0.537. `Judge_label` is the LLM-as-judge verdict, *Yes* or not-*Yes*, by majority vote over five runs. `Truncated` indicates whether generation reached the 128-token cap without emitting an end-of-sequence token.

#### Curation

`Curator_score` is the functional similarity rating, on the following scale:

- 5, *equivalent* : the same function and role, wording may differ;
- 4, *highly similar* : the same core function under a different emphasis, with no conflicting claims;
- 3, *related* : the same general functional category but distinct specific roles;
- 2, *somewhat related* : only a broad high-level connection, no agreement on the actual function;
- 1, *unrelated or contradictory*.

`Specificity` and `Information` give the direction of each relative to the Swiss-Prot description. `SwissProt_description` and `ProtTale_prediction` are the two texts the curator compared, and Curator notes is the free-text justification, present for 180 of the 232 items.

Neither `p_JAY` nor `Judge_label` was visible to the curator at scoring time; both are included here so that the human ratings can be compared against the automated ones directly.

### Supplementary Data

All Supplementary Data are at https://huggingface.co/Mulah/ProtTale/tree/main/data, where a README defines every column.

**Supplementary Data 1** (`Supplementary_Data_1_expert_curation.csv`). The 232 expert-rated predictions (Supplementary Note 8).

**Supplementary Data 2** (`Supplementary_Data_2_unseen_per_method.csv`). All 1,031 unseen proteins for each of the five methods and the curated-text reference: the curated Swiss-Prot description and each method’s generated description, the judge label from each of five runs and the majority, the extracted GO terms and Wang similarity from each extraction run, taxonomic group, length, and ProtTale’s reliability score with whether it passes the discrete filter, which 201 of the 1,031 proteins do. Source data for Figs. 1–4 and Extended Data Figs. 3B and 4.

**Supplementary Data 3** (`Supplementary_Data_3_stage1_test_pairs.csv`). The 534 Stage 1 test pairs with their MIPS label and the judge label from a single run (Extended Data Fig. 3A).

**Supplementary Data 4** (`Supplementary_Data_4_shuffled_baseline.csv`). Reliability scores for real and residue-shuffled sequences (Extended Data Fig. 5).

**Supplementary Data 5** (`Supplementary_Data_5_mips_validation.xlsx`). The 1,000 same-cluster pairs used to validate MIPS (Supplementary Note 1).

**Supplementary Data 6** (`Supplementary_Data_6_training_curves.csv`). Training and validation curves for both stages and the reliability head (Extended Data Fig. 9).

**Supplementary Data 7** (`Supplementary_Data_7_ablation_no_stage1.csv`). Predictions and judge labels of ProtTale w/o Stage 1 (Extended Data Fig. 7).

Over the unseen set, 11% of proteins score below 0.20, 28% fall in 0.20–0.30, 57% in the 0.30–0.70 band that expert curation did not sample, and 3% reach 0.70 or above.

